# Placental and fetal microbiota in rhesus macaque: a case study using metagenomic sequencing

**DOI:** 10.1101/2024.08.16.608220

**Authors:** Qiao Du, Xu Liu, Rusong Zhang, Gang Hu, Qinghua Liu, Wen Ma, Ying Hu, Zhenxin Fan, Jing Li

**Affiliations:** Key Laboratory of Bio-Resources and Eco-Environment (Ministry of Education), College of Life Sciences, Sichuan University, Chengdu, Sichuan, China; SCU-SGHB Joint Laboratory on Non-human Primates Research, Sichuan Green-house Biotech Co., Ltd., Meishan, Sichuan, China

**Keywords:** metagenomics, rhesus macaques, NHP, fetal microbiota, immunity

## Abstract

Recent evidence challenging the notion of a sterile intrauterine environment has sparked research into the origins and effects of fetal microbiota on immunity development during gestation. Rhesus macaques (RMs) serve as valuable non-human primate (NHP) models due to their similarities to humans in development, placental structure, and immune response. In this study, metagenomic analysis was applied to the placenta, umbilical cord, spleen, gastrointestinal (GI) tissues of an unborn RM fetus, and the maternal intestine, revealing the diversity and functionality of microbes in these tissues. We observed substantial microbial sharing between the mother and fetus, with the microbial composition of the placenta and umbilical cord more closely resembling that of the fetal organs than the maternal intestine. Notably, compared with other adult RMs, there was a clear convergence between maternal and fetal microbiota, alongside distinct differences between the microbiota of adults and the fetus, which underscores the unique microbial profiles in fetal environments. Furthermore, the fetal microbiota displayed a less developed carbohydrate metabolism capacity than adult RMs. It also shared antibiotic resistance genes (ARGs) with both maternal and adult RM microbiomes, indicating potential vertical transmission. Comparative analysis of the metagenomes between the RM fetus and a human fetus revealed significant differences in microbial composition and genes, yet also showed similarities in certain abundant microbiota. Collectively, our results contribute to a more comprehensive understanding of the intrauterine microbial environment in macaques.

## 1 Introduction

Activation of the fetal immune system during gestation significantly impacts fetal health and pregnancy outcomes (1–8). A well-developed immune system with a diverse immune repertoire is essential for the fetus to respond effectively to potential antigens and other threats (6, 9, 10). Fetal inflammatory response, closely linked to immune system activation, is a key indicator of overall fetal health (6, 7, 11, 12). Additionally, diverse immune cells present in the fetal gut may initiate immune activation, highlighting the gut as a critical site for immune responses (13–17). Initial studies predominantly supported the sterile womb hypothesis, suggesting that the placenta functions as an immune organ protecting the fetus from exposure to external antigens and microbes during pregnancy (18–20). Furthermore, the placenta’s role extends beyond mere barrier protection, encompassing maternal-fetal cell communication and regulation of immune interactions (21–23).

However, an increasing number of studies provided evidence of low-biomass microbes in human placental and fetal tissues (24–31). Notably, early exposure to microbes can influence the developing immune system, subsequently affecting reproduction and pregnancy outcomes (32–36). Amid concerns about potential contamination, robust research methods, including the use of negative controls, culture techniques, and scanning electron microscopy, have successfully validated the presence of microorganisms in the intrauterine environment (35, 37–40). These studies challenged prior beliefs, reshaping our understanding of microbial vertical transmission and its role in developing immune system and reproductive outcomes.

The primary limitation in current research on human placental and fetal microbiota is the difficulty in obtaining placental and fetal tissue before delivery. Post-delivery tissue collection introduces confounding variables that can potentially skew experimental results (29, 41–43). To address these challenges, researchers have increasingly turned to animal models, with non-human primates (NHPs) proving particularly valuable due to their close physiological, anatomical, reproductive, genetic, behavioral, and immune similarities to humans (44–48). Among NHPs, Rhesus macaques (RMs), noted for their developmental patterns, placental structures, and immune responses highly similar to those of humans, are especially significant, making them an effective model for studying microbial infection and fetal immunity in pregnancy (47–52). However, research on intrauterine microbiota in RMs remains relatively scarce, largely due to ethical and welfare concerns associated with collecting fetal samples from NHPs (53, 54). Several studies using 16S rRNA sequencing to investigate microbial signals in fetal and placental tissue of RMs and Japanese macaques have reached controversial conclusions regarding the presence of intrauterine microbiota (55–60). Given these challenges and the existing controversies, further targeted research is essential.

In the current study, we utilized cesarean section to collect fetal samples from a deceased pregnant RM and conducted a comprehensive metagenomic analysis of fetal microbes. Despite the implementation of stringent criteria to exclude environmental contamination and low-quality sequences, we observed a variety of metabolically active microbes in the placenta, umbilical cord, and fetal organs. To explore potential vertical transmission in RMs, we compared the microbial profiles of the mother and the fetus. Comparisons of microbial composition, function, and antibiotic resistance genes (ARGs) were conducted between fetal, maternal, and other adult RMs provided further evidence supporting the possibility of vertical transmission. Separately, we compared the microbiota between the RM fetus and a human fetus, revealing key differences and similarities and suggesting the potential of RMs as a model for studying human fetal microbiomes. This study aims to contribute valuable information to the understanding and research of fetal microbiota and its potential impact on fetal immunity and overall health.

## 2 Materials and Methods

### 2.1 Sample collection

Maternal intestinal contents, as well as placental, umbilical cord, fetal spleen, and fetal gastrointestinal (GI) tissues, were obtained from a captive female RM housed at the Sichuan Green-House Biotech Co., Ltd. (Meishan, Sichuan, China). The company was established by West China Hospital of Sichuan University and has gained the License for Experimental Animals and Production of Experimental Animal Feed and License for Use of Experimental Animals. All the macaques are conducted periodical physical examinations by professional veterinaries and there is a daily inspection of all the monkey rooms. The dietary composition of the macaque in this experiment was as follows: 50% corn, 14% wheat, 15% soybean meal, 5% wheat bran, 4.5% fish meal, 5.5% sucrose, 5% bone meal, 0.1% multivitamins, 0.1% vitamin C powder, 0.1% trace elements, 0.1% lysine, 0.2% methionine, and 0.1% vitamin B complex powder. All procedures were conducted in accordance with and under the approval of the Ethics Committee of the College of Life Sciences, Sichuan University, China.

The subject of our study was an eight-year-old maternal macaque. The macaque, in the late stage of pregnancy, died around 19th to 20th weeks of gestation. The maternal macaque was under the vigilant care of seasoned caregivers and was kept under long-term observation. Notably, no evident signs of illness were observed in the macaque during both the pregnancy and pre-pregnancy periods. A qualified veterinarian conducted an autopsy on the maternal macaque less than two hours after its death, revealing no obvious pathological anomalies in the tissues. The fetus’s death was a consequence of the mother’s death. According to the caregivers’ records, the maternal macaque died on August 10, 2022. The death of the maternal macaque was probably caused by the stress from high temperatures during that period.

Before the dissection of the maternal macaque, the entire ultraclean workbench was disinfected thoroughly. Intestinal contents from the mother were collected using aseptic equipment. The fetus, which did not pass through the birth canal, was retrieved after excision of the entire sterilized uterus, which was placed in an ultraclean workbench for ultraviolet irradiation. All samples were collected using sterile instruments. Before dissection, swabs of the workbench and operator hands were used as environmental controls, samples of phosphate-buffered saline (PBS) wash of the dissection instruments were used as PBS controls, and blank swabs and PBS solution were used as blank controls (Figure S1). Although we did not perform metagenomic sequencing on these control samples due to their low DNA content, we conducted library preparation for DNA quantification to verify that the potential contribution of environmental contamination to our sample data was minimal (mean ± SD ng/μL, controls: 0.48 ± 0.37).

### 2.2 Metagenomic sequencing and quality control

All DNA samples were extracted using a Tiangen DNA Stool Mini Kit (Tiangen Biotech Co., Ltd., China) and sent to Novogene Bioinformatics Technology Co., Ltd. (Beijing, China) for metagenomic sequencing. In total, 0.2 μg of DNA per sample was used as input for the DNA library preparations, and the sequencing library was generated using a NEBNext® UltraTM DNA Library Prep Kit for Illumina (NEB, USA, Catalog #: E7370L). After library quality assessment and quantification, the qualified libraries were sequenced using the Illumina 6000 platform with a paired-end length of 150 bp. The Q30 and Q20 values of all samples were approximately 90% and 95%, respectively, indicating that the correct rate of base calling was relatively high (Table S1). Adapters and low-quality reads in the raw data were removed using Fastp (61). Host contamination was removed using Bowtie2 (62) based on the RM reference genome (assembly Mmul_10).

Additionally, gut metagenomic data of seven healthy adult RMs were included for comparison with fetuses, two of which were downloaded from the China National GeneBank Database (CNGBdb) (accession number CNP0002963), and the remaining five were obtained from our laboratory. We downloaded three metagenomic datasets of intestinal contents from a single human fetus, sourced from the Genome Sequence Archive for Human (accession number HRA003676). These samples were collected using aseptic techniques from elective pregnancy terminations during the 19th to 20th weeks.

### 2.3 Data processing and analyses

Metagenome *de novo* assembly was conducted using MEGAHIT (63) with the option “–m 0.95 -- min-contig-len 300”. The genes in the metagenomic data were obtained using Prodigal (64) with the option “-p meta”. Non-redundant genes were constructed using CD-HIT (65) with the option “-c 0.95 -aS 0.90” and quantified using Salmon (66) with the option “--meta”. The non-redundant amino acid sequences translated from the genes were used for subsequent functional prediction. Functional annotations of carbohydrate-active enzymes (CAZymes) and ARGs were performed using DIAMOND (67) with the option “--id 80% --query-cover 70% --evalue 1e-5” and rgi based on the Carbohydrate Active enZYmes database (68) and comprehensive antibiotic resistance database (69). Total abundance of each functional gene type was the sum of the abundances of all genes mapped to the same type. The abundances of CAZyme genes and ARGs were normalized by transcripts per million (TPM). The microbial metabolic pathways and gene families were predicted using HUMANn3 (70) based on the ChocoPhlAn and UniRef90 EC filtered databases (71), and abundances were normalized by counts per million (CPM). The taxonomic annotation of metagenomic sequences was performed using kraken2 (72) with the option “--use-mpa-style”. The abundances of taxa were normalized to relative abundance.

Principal coordinate analysis (PCoA) and Bray-Curtis distance were calculated using the Adonis test in the vegan package of the R statistical environment (version 4.2.3) (73). Alpha-diversity analysis was done by the vegan package of R statistical software with Wilcoxon’s rank-sum test. Statistical significance of differences in numbers were analyzed using the Wilcoxon’s rank-sum test with *p* < 0.05. Differentially abundant pathways were screened by STAMP (v2.1.3) based on Welch’s t-test (FDR < 0.05; Benjamini–Hochberg method).

## 3 Results

### 3.1 Microbial composition of maternal intestine, placenta, umbilical cord, and fetal organs

Microbial signals in the cesarean-delivered RM fetus were investigated. To minimize background contamination, environmental, PBS, and blank controls were utilized, with undetectable (below sequencing thresholds) microbial DNA indicating negligible impact on the results. Metagenomic sequencing of maternal intestinal content and placental, umbilical cord, fetal spleen, and fetal GI tissues identified 2 806 species of bacteria (belonging to 31 phyla and 891 genera, Figure 1A-C), 242 species of viruses (belonging to eight orders and 102 genera, Figure 2A-C), and 74 species of archaea (belonging to three phyla and 44 genera, Figure 3A-C). A PCoA plot, based on Bray-Curtis distances of microbial species-level relative abundance profiles, revealed that samples from the placenta, umbilical cord, fetal spleen, and fetal GI clustered closely together, distinct from the maternal intestinal sample (Figure S2A). For comparative analysis of microbial composition between fetal and maternal samples, samples from the placenta, umbilical cord, fetal spleen, and fetal GI were grouped as the “fetal group” for further analysis. Within this fetal group, a total of 2 554 species of bacteria (belonging to 28 phyla and 826 genera), 223 species of viruses (belonging to eight orders and 97 genera), and 68 species of archaea (belonging to three phyla and 142 genera) were identified.

**Figure 1.**
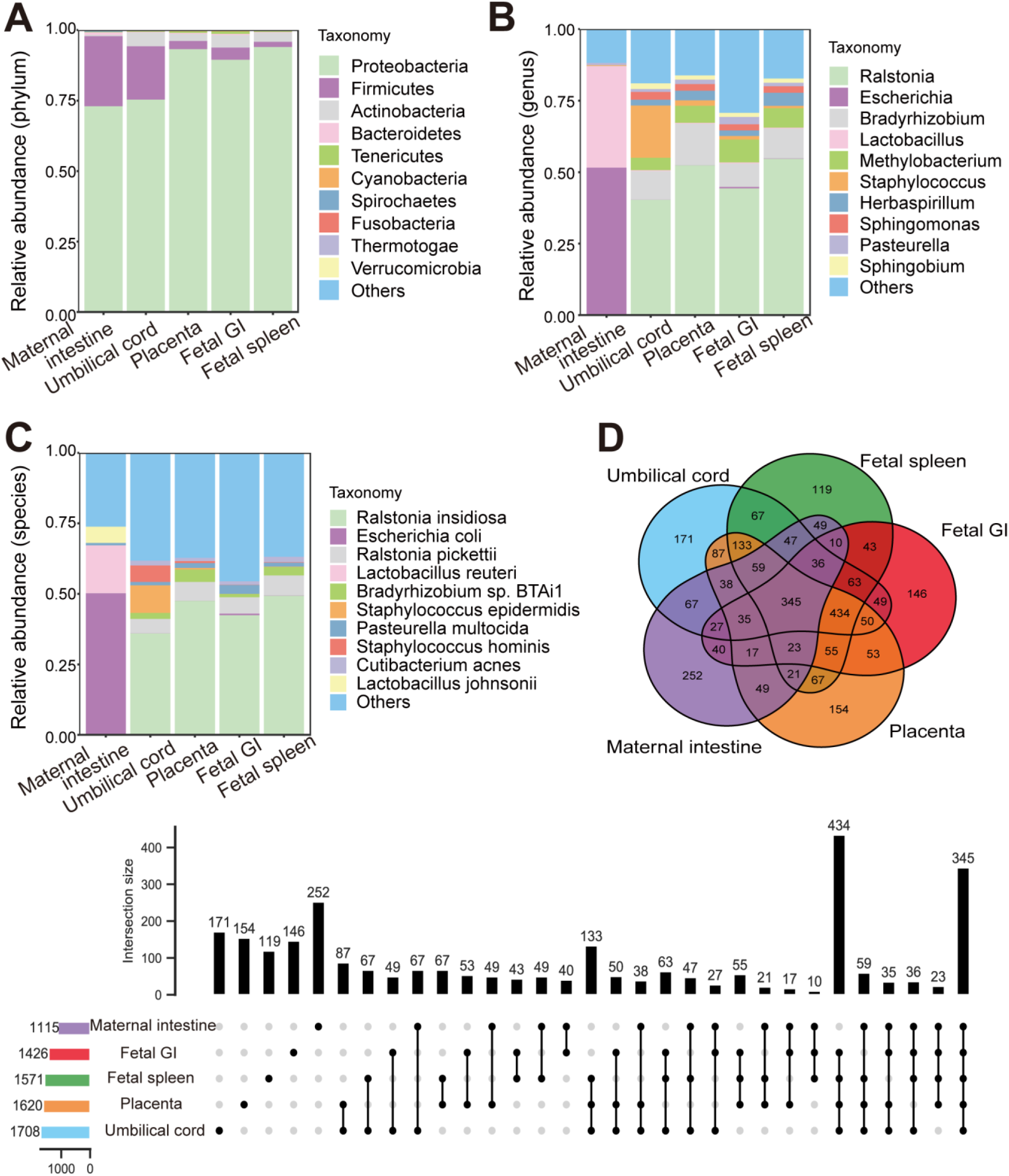
Bacterial distribution in maternal and fetal samples. (**A**) Main bacteria phyla with the highest relative abundance in different samples. (**B**) Main bacteria genera with the highest relative abundance in different samples. (**C**) Main bacteria species with the highest relative abundance in different samples. (**D**) UpSet diagram and Venn diagram of exclusive and shared maternal and fetal bacteria in different samples.

**Figure 2.**
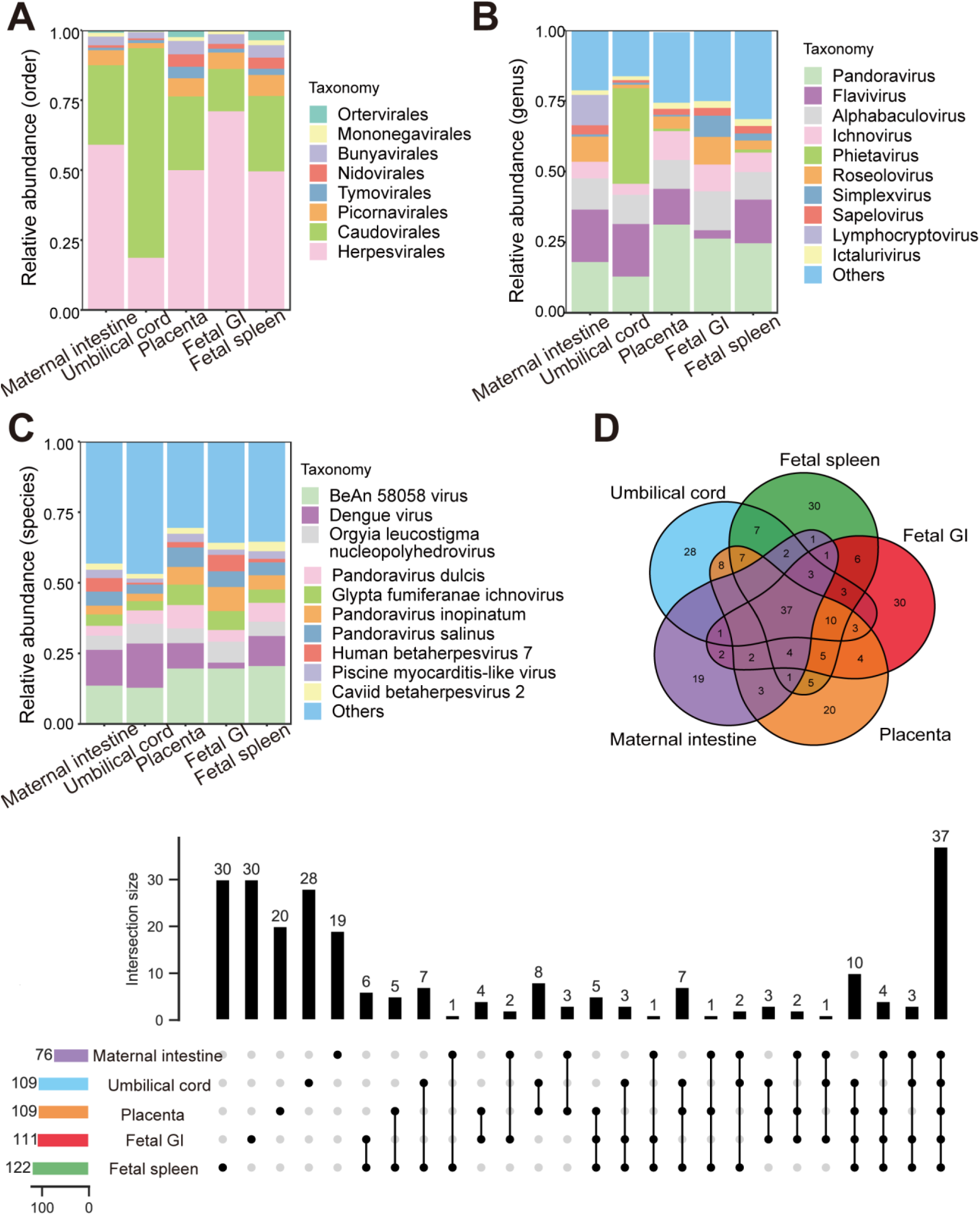
Viral distribution in maternal and fetal samples. (**A**) Main virus orders with the highest relative abundance in different samples. (**B**) Main virus genera with the highest relative abundance in different samples. (**C**) Main virus species with the highest relative abundance in different samples. (**D**) UpSet diagram and Venn diagram of exclusive and shared maternal and fetal viruses in different samples.

**Figure 3.**
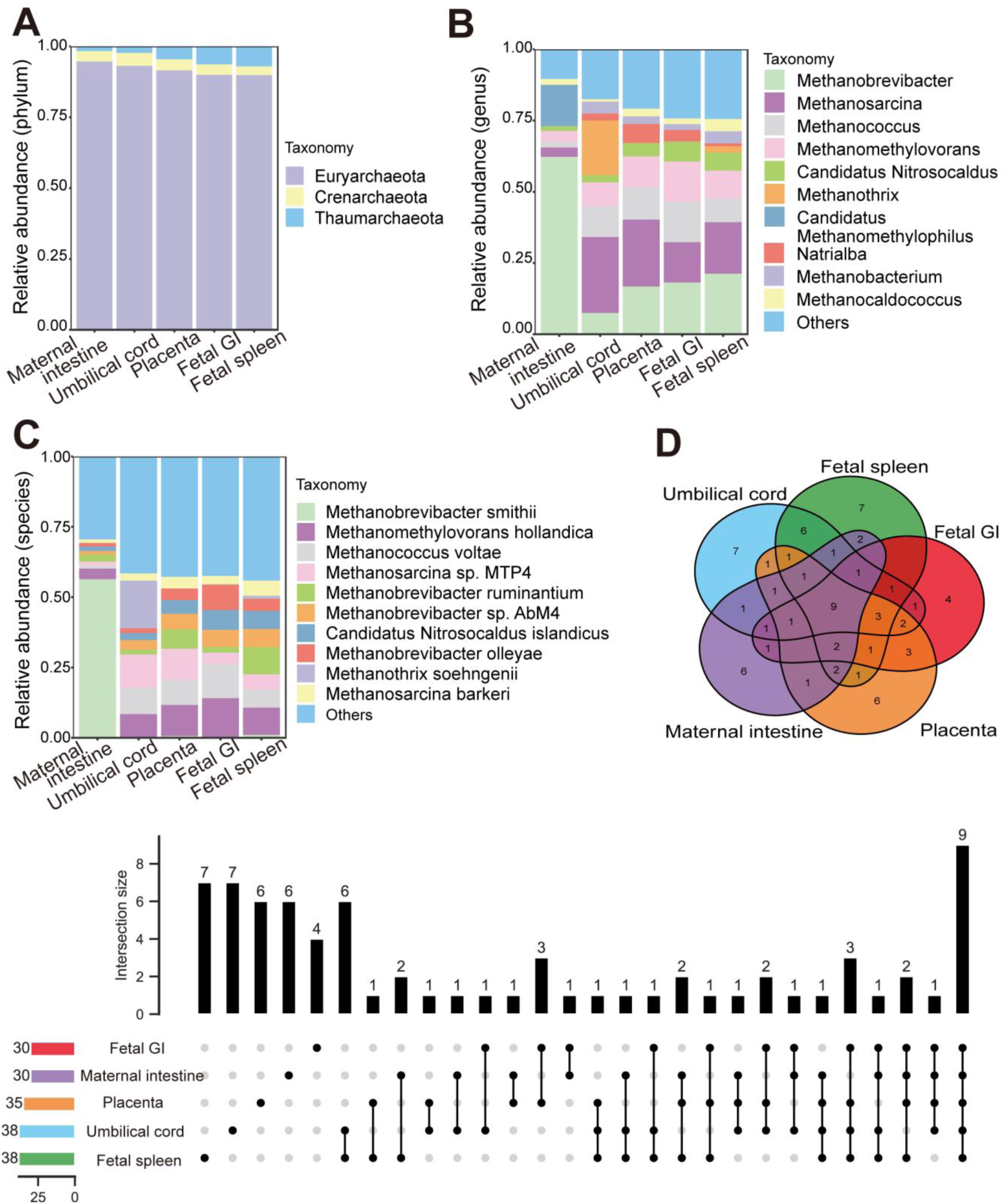
Archaeal distribution in maternal and fetal samples. (**A**) Main archaea phyla with the highest relative abundance in different samples. (**B**) Main archaea genera with the highest relative abundance in different samples. (**C**) Main archaea species with the highest relative abundance in different samples. (**D**) UpSet diagram and Venn diagram of exclusive and shared maternal and fetal archaea in different samples.

Among the 2 806 bacterial species identified, 345 were shared by the maternal and fetal groups, with the number of organ-unique bacteria varying from 119 (fetal spleen) to 252 (maternal intestine) (Figure 1D). In total, 779 out of 2 554 bacterial species were shared by the placenta, umbilical cord, fetal spleen, and fetal GI samples, each containing 203, 238, 168, and 186 unique bacteria species (Figure S2B). Notably, each fetal sample shared considerable bacterial species with the maternal intestinal sample, varying from 533 to 654 species (Table S2). Among the most dominant bacteria in the maternal and fetal samples (Table 1), Proteobacteria and Firmicutes were the dominant shared phyla in both groups. Interestingly, no dominant shared genera or species were found between the maternal and fetal groups. Within the fetal group, the dominant bacteria in the placenta, umbilical cord, fetal GI, and spleen samples were generally identical, although *Staphylococcus* was a dominant genus in the umbilical cord but not in the other fetal samples.

**Table 1.** Top five bacteria with the highest relative abundance in maternal and fetal RMs.

| Level | Maternal intestine | Placenta | Umbilical cord | Fetal GI tissue | Fetal spleen |
| --- | --- | --- | --- | --- | --- |
| Phylum | Proteobacteria | Proteobacteria | Proteobacteria | Proteobacteria | Proteobacteria |
|  | Firmicutes | Firmicutes | Firmicutes | Actinobacteria | Actinobacteria |
|  | Bacteroidetes | Actinobacteria | Actinobacteria | Firmicutes | Firmicutes |
|  | Spirochaetes | Tenericutes | Bacteroidetes | Tenericutes | Tenericutes |
|  | Actinobacteria | Bacteroidetes | Tenericutes | Bacteroidetes | Bacteroidetes |
| Genus | *Escherichia* | *Ralstonia* | *Ralstonia* | *Ralstonia* | *Ralstonia* |
|  | *Lactobacillus* | *Bradyrhizobium* | *Staphylococcus* | *Bradyrhizobium* | *Bradyrhizobium* |
|  | *Campylobacter* | *Methylobacterium* | *Bradyrhizobium* | *Methylobacterium* | *Methylobacterium* |
|  | *Streptococcus* | *Herbaspirillum* | *Methylobacterium* | *Enterobacter* | *Herbaspirillum* |
|  | *Shigella* | *Sphingomonas* | *Sphingomonas* | *Pasteurella* | *Sphingomonas* |
| **Species** | *Escherichia coli* | *Ralstonia insidiosa* | *Ralstonia insidiosa* | *Ralstonia insidiosa* | *Ralstonia insidiosa* |
|  | *Lactobacillus reuteri* | *Ralstonia pickettii* | *Staphylococcus epidermidis* | *Ralstonia pickettii* | *Ralstonia pickettii* |
|  | *Lactobacillus johnsonii* | *Bradyrhizobium sp. BTAi1* | *Staphylococcus hominis* | *Pasteurella multocida* | *Bradyrhizobium sp. BTAi1* |
|  | *Lactobacillus amylovorus* | *Pasteurella multocida* | *Ralstonia pickettii* | *Enterobacter cloacae* | *Herbaspirillum seropedicae* |
|  | *Campylobacter hyointestinalis* | *Bradyrhizobium diazoefficiens* | *Bradyrhizobium sp. BTAi1* | *Cutibacterium acnes* | *Cutibacterium acnes* |

Compared with the identified bacteria, the proportion of identified viruses and archaea was notably small, with only 0.33%-1.85% viruses and 0.097%-0.18% archaea, respectively (Figure S2E). Among the 242 virus species identified, 37 were shared by the maternal intestine and fetal group, each sample harboring 19–30 organ-specific viruses (Figure 2D). The four fetal samples (placenta, umbilical cord, fetal spleen, and fetal GI) shared 47 virus species, each harboring 23, 28, 31, and 32 unique viruses, respectively (Figure S2C). And there were 43-50 virus species were share by the individual fetal samples and maternal intestinal sample (Table S2). Among the most abundant viruses, Caudovirales, Herpesvirales, and Picornavirales were the dominant shared orders between the maternal and fetal groups (Table 2), and *Pandoravirus* and *Alphabaculovirus* were the dominant shared genera. Most of the dominant virus genera were consistent across the four fetal samples, although the umbilical cord sample differed somewhat from the other fetal samples. Among the 74 archaea species identified, nine were common between the maternal and the fetal groups, with each harboring 4–7 unique species (Figure 3D). In total, 12 archaeal species were shared by the four fetal samples, with each fetal sample (Figure S2D). Furthermore, 15–18 archaeal species were shared by the maternal and four fetal samples (Table S2). Both groups contained the same dominant phyla (Table 3), although the umbilical cord sample differed somewhat in dominant genera.

**Table 2.** Top five viruses with the highest relative abundance in maternal and fetal RMs.

| **Level** | **Maternal intestine** | **Placenta** | **Umbilical cord** | **Fetal GI tissue** | **Fetal spleen** |
| --- | --- | --- | --- | --- | --- |
| **Order** | Herpesvirales | Herpesvirales | Caudovirales | Herpesvirales | Herpesvirales |
|  | Caudovirales | Caudovirales | Herpesvirales | Caudovirales | Caudovirales |
|  | Picornavirales | Picornavirales | Bunyavirales | Picornavirales | Picornavirales |
|  | Bunyavirales | Bunyavirales | Picornavirales | Bunyavirales | Bunyavirales |
|  | Mononegavirales | Nidovirales | Tymovirales | Nidovirales | Nidovirales |
| **Genus** | *Flavivirus* | *Pandoravirus* | *Phietavirus* | *Pandoravirus* | *Pandoravirus* |
|  | *Pandoravirus* | *Flavivirus* | *Flavivirus* | *Alphabaculovirus* | *Flavivirus* |
|  | *Alphabaculovirus* | *Alphabaculovirus* | *Pandoravirus* | *Roseolovirus* | *Alphabaculovirus* |
|  | *Lymphocryptovirus* | *Ichnovirus* | *Alphabaculovirus* | *Ichnovirus* | *Ichnovirus* |
|  | *Roseolovirus* | *Roseolovirus* | *Ichnovirus* | *Simplexvirus* | *Roseolovirus* |
| **Species** | *BeAn 58058 virus* | *BeAn 58058 virus* | *Dengue virus* | *BeAn 58058 virus* | *BeAn 58058 virus* |
|  | *Dengue virus* | *Dengue virus* | *BeAn 58058 virus* | *Pandoravirus inopinatum* | *Dengue virus* |
|  | *Macacine gammaherpesvirus 4* | *Pandoravirus dulcis* | *Orgyia leucostigma nucleopolyhedrovirus* | *Orgyia leucostigma nucleopolyhedrovirus* | *Pandoravirus dulcis* |
|  | *Pandoravirus salinus* | *Glypta fumiferanae ichnovirus* | *Staphylococcus phage StB27* | *Glypta fumiferanae ichnovirus* | *Pandoravirus inopinatum* |
|  | *Orgyia leucostigma nucleopolyhedrovirus* | *Pandoravirus salinus* | *Staphylococcus virus PH15* | *Human betaherpesvirus 7* | *Orgyia leucostigma nucleopolyhedrovirus* |

**Table 3.**
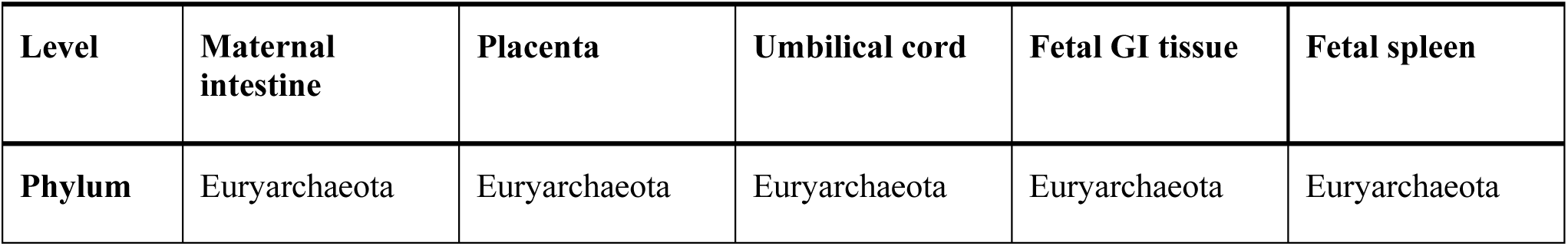
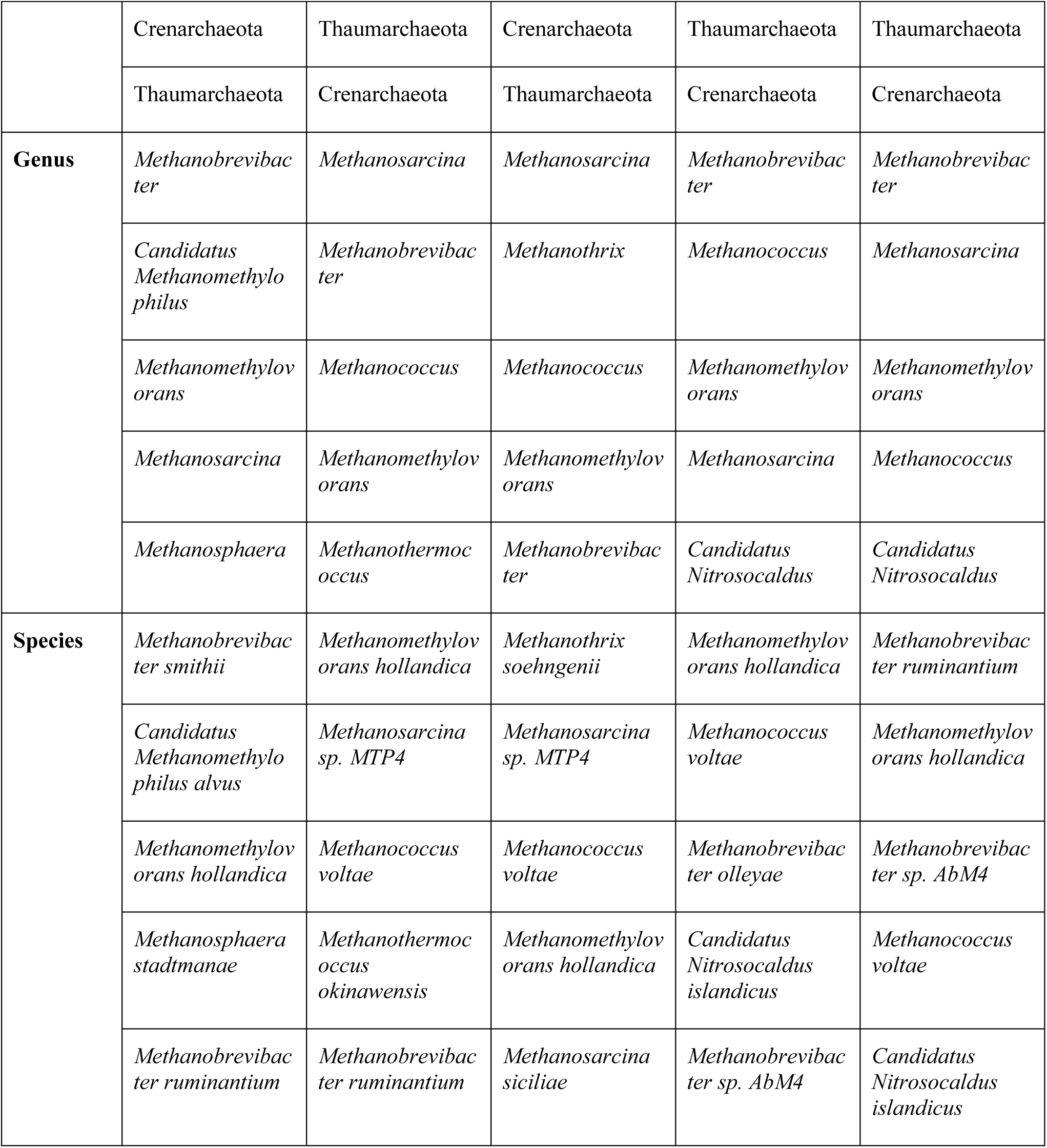
Top five archaea with the highest relative abundance in maternal and fetal RMs.

### 3.2 Comparison of microbial composition and function between fetal and adult RMs

To assess microbial differences between fetal and adult RMs, seven gut metagenomic datasets obtained from adult RMs were included for analysis. Comparing α-diversity between the adult and fetal groups revealed significant differences in the ACE and Chao1 indices (richness) (*p* < 0.05; Figure 4A), but no significant differences in the Shannon and Simpson indices (richness and heterogeneity). Notably, the number of microbial species was significantly higher in adult samples compared to the fetal samples (*p* < 0.05; Figure 4B). The heatmap of microbes at the genus level indicated that placental, umbilical cord, fetal spleen, and fetal GI samples clustered separately from maternal intestinal and other adult RM samples, highlighting the distinct microbial composition between the adult and fetal groups (Figure 4C). Abundant microbes exhibited differences between the adult and fetal groups. Specifically, the predominant bacteria in the adult microbiome were Bacteroidetes and Firmicutes, whereas Proteobacteria was dominant in the fetal microbiome (Table S3). It is evident that while the maternal samples clustered with other adult RM samples, their microbial composition closely aligned with that of the fetal group (Figure 4C). The PCoA results based on microbial species-level abundance further corroborated the heatmap results (Figure 4D). Furthermore, the PCoA results for viruses, bacteria, and archaea (Figure S3) showed that viral composition between the fetal and maternal samples was more similar (Figure S3A) than their bacterial or archaeal compositions (Figure S3B, C).

**Figure 4.**
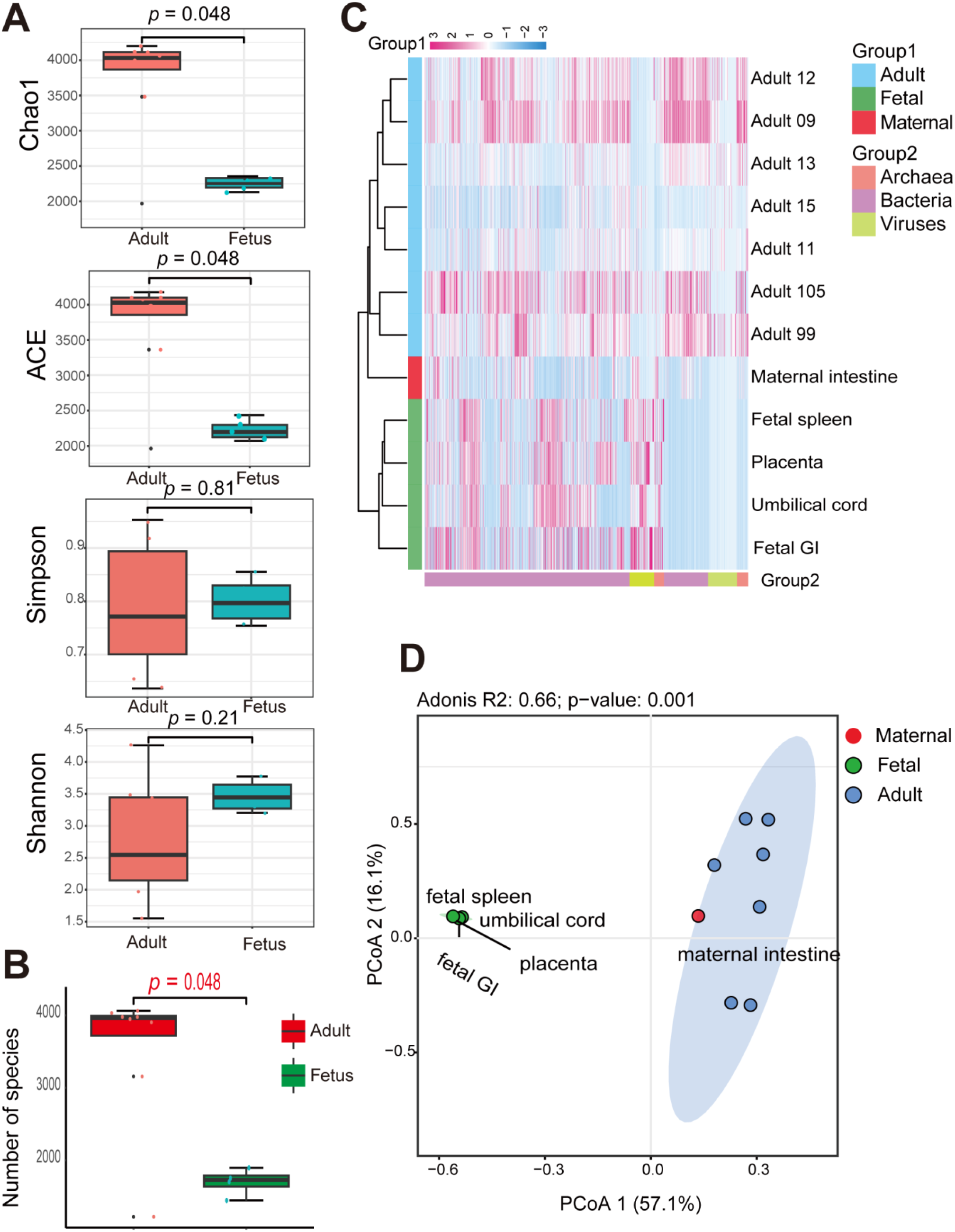
Different analyses of microbial composition between adult and fetal RMs. (**A**) Alpha diversity estimates (ACE, Chao1, Shannon, and Simpson indices) between fetal and adult groups (**B**) Comparison of microbial species number in adult and fetal groups (Wilcoxon’s rank-sum test, *p* < 0.05). (**C**) Heatmap plot of bacteria, archaea, and viruses in each sample at genus level. (**D**) PCoA plot based on Bray-Curtis distance of species-level relative abundance of microbiota between adult and fetal samples (Adonis, R^2^ = 0.66, *p* = 0.001).

Functional differences between the fetal and adult groups were also studied. The PCoA analysis demonstrated a distinct separation between fetal and adult microbial gene families, with maternal samples displaying a closer similarity to those of the fetal microbiomes (Figure S3D). Building on these genetic findings, we next explored the associated metabolic pathways. The microbiota of the adult group was primarily enriched in protein, ribonucleotide, and peptidoglycan synthesis pathways (Figure S4A), while the fetal group was primarily enriched in pathways related to nucleotide and amino acid synthesis, fatty acid synthesis and oxidation, and energy metabolism (Figure S4B-E). A total of 39 significantly distinct metabolic pathways were identified between the groups (Figure 5A). Of these, 28 pathways were enriched in the adult group, which were mainly related to carbohydrate metabolism essential for energy conversion, metabolic regulation, and energy balance. Conversely, of the 11 pathways enriched in the fetal group, the most significant were associated with energy synthesis and metabolism, including molybdopterin biosynthesis, aerobic respiration I (cytochrome c), and heme b biosynthesis II (oxygen-independent).

**Figure 5.**
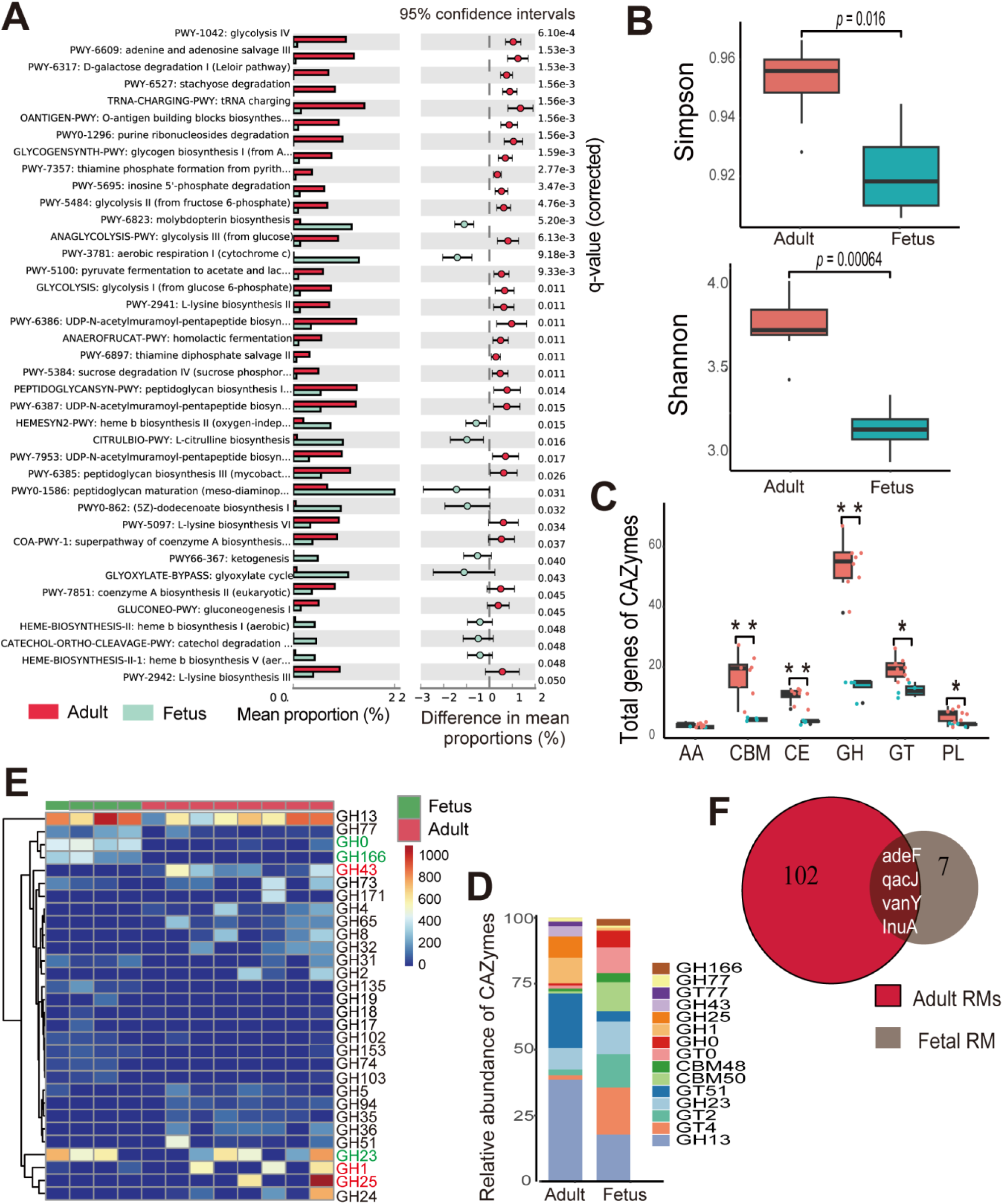
Functional comparison of microbiota between adult and fetal RMs. (**A**) Functionally predicted pathways differing in abundance in adult and fetal groups. Bar plot shows mean proportions of pathways. Only *p* < 0.05 is shown (Welch’s *t*-test, false discovery rate (FDR)-adjusted). (**B**) Alpha diversity estimates (Shannon and Simpson indices) of CAZyme genes between fetal and adult groups. (**C**) Gene number comparisons of GH, GT, CBM, CE, PL, and AA. \**p*-value < 0.05, \*\**p*-value < 0.01 (**D**) Main CAZymes with highest relative abundance in adult and fetal microbiome. (**E**) Heatmap plot of abundance of top 30 GH family genes. GHs marked in red or green are most abundant in RM or human fetal groups. Red GHs mean significantly more abundant in adult microbiome of RMs, and green GHs mean significantly more abundant in fetal microbiome. (**F**) Distribution of ARGs in two groups.

We next identified microbial CAZymes, including glycoside hydrolase (GH), glycosyl transferase (GT), carbohydrate-binding module (CBM), carbohydrate esterase (CE), polysaccharide lyase (PL), and auxiliary activity (AA) between the two groups. The Shannon and Simpson indices indicated that CAZyme diversity was significantly higher in the adult group than in the fetal group (*p* < 0.05; Figure 5B). The adult microbiome contained a significantly greater number of CAZyme genes than the fetal microbiome (Figure S4F). Notably, the adult samples displayed significantly more GHs, GTs, CBMs, CEs, and PLs than the fetal samples (Figure 5C). In both groups, GH was the most abundant family, followed by GT. The top five prevalent CAZyme families in the adult group were GH13, GT51, GH1, GH25, and GH23, and in the fetal group were GT4, GH13, GT2, GH23, and CBM50 (Figure 5D). Given the critical role of the GH family in carbohydrate degradation, we compared the abundances of different GH family members between the adult and fetus groups. GH13 was the most abundant GH family member, while GH1, GH25, and GH43 were more abundant in the adult group, and GH23, GH0, and GH166 were more abundant in the fetal group (Figure 5E and S4G). Many GH family genes abundant in the adult group were less so in the fetal group. Notably, seven ARGs were identified in the fetal samples (six ARGs in umbilical cord, one ARG in placenta, one ARG in fetal spleen, and two ARGs in fetal GI) and 102 ARGs were identified in the adult group, including the mother. Among them, four ARGs (adeF, qacJ, vanY, and InuA) were shared by the adult and fetal groups (Figure 5F).

### 3.3 Comparison of fetal microbial composition between the RM and human

Metagenomic datasets of intestinal contents from a single human fetus were downloaded for comparison with the RM fetus data. A total of 96 species of viruses (belonging to 52 genera), 32 species of archaea (belonging to three phyla and 19 genera), and 982 species of bacteria (belonging to 20 phyla and 431 genera) were identified in the human fetal samples. Comparison of α-diversity between the two species showed no significant differences in Shannon and Simpson indices (Figure 6A). However, at microbial species-level abundance, PCoA distinctly separated the human and RM fetal samples (*p* < 0.05; Figure 6B). A similar distinction emerged following PCoA of gene family abundance of microbiota (*p* < 0.05, Figure 6C).

**Figure 6.**
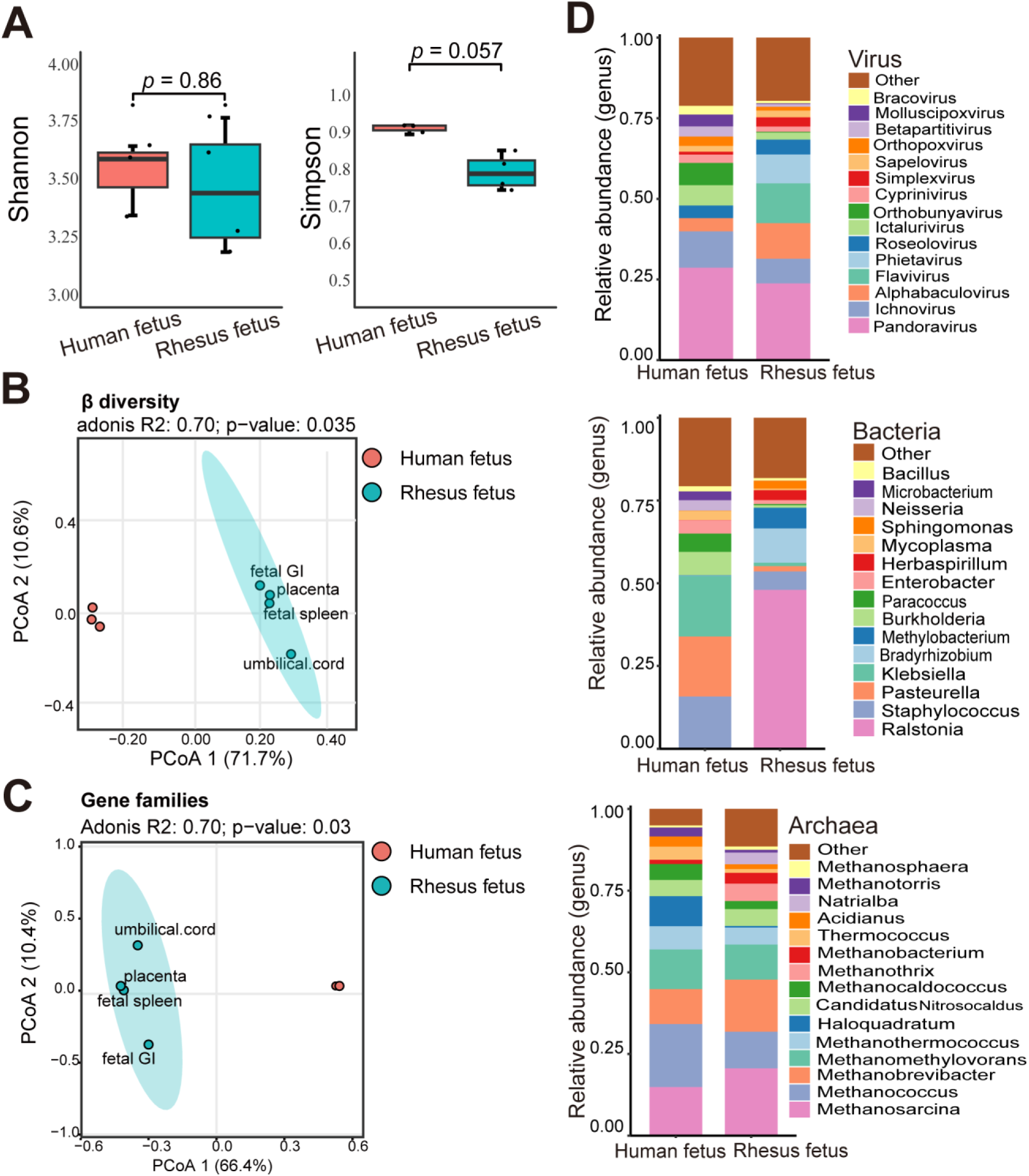
Different analysis of microbial composition between human and RM fetal samples. (**A**) Alpha diversity estimates (Shannon and Simpson indices) between two groups (Wilcoxon’s rank-sum test, *p* > 0.05). (**B**) PCoA plot based on Bray-Curtis distance of species-level relative abundance of microbiota between human and RM fetal samples (Adonis, R^2^ = 0.7, *p* = 0.034). (**C**) PCoA plot based on gene family abundance between human and RM fetal samples (Adonis, R^2^ = 0.66, *p* = 0.031). (**D**) Top 15 abundant microbial genera (virus, archaea, and bacteria) in two groups. Comparing microbial composition between the RM and human fetal samples, bacteria exhibited more pronounced differences at the genus levels than viruses and archaea (Figure 6D). Despite the differences in microbiota between the two species, several microbes were shared. For instance, both the RM and human fetal samples contained methanogens being the dominant archaea (Table S4). The dominant bacterial phyla in the fetal microbiota of both species displayed a high degree of similarity (Figure S5 and Table S5).

## 4 Discussion

### 4.1 Microbes identified in RM fetus

Previous studies indicate that the development of the human immune system begins early in fetal life (5, 6, 35), with exposure to maternal antigens and microbial entities that traverse the placental barrier into fetal tissues (13, 33, 37, 74). This microbial and antigenic exposure is believed to prime the fetal immune system, establishing a foundation for lifelong immunity and tolerance (35). As a widely used animal model, RMs offer a valuable opportunity to investigate the fetal microbiota (47–52). In the present study, we performed metagenomic analysis on fetal organs obtained in a sterile manner by cesarean section, and further collected samples of placenta, umbilical cord, and maternal intestine content to study microbial signal in fetus. Following rigorous experimental design, including the exclusion of background contamination and removal of low-quality and host sequences, we detected diverse microbes (including 223 virus species, 2 554 bacteria species, and 68 archaea species) in the placental, umbilical cord, and fetal organ samples. A considerable number of shared microbes were found in the placenta, umbilical cord, and fetal organs, with certain microbes exhibiting significant similarity to those in the maternal intestine. Our findings suggest that in RMs, the fetus is exposed to maternal microbes prior to birth due to vertical transmission. These findings are consistent with previous studies in humans, NHPs, and mice that show the presence of microbes in the intrauterine environment (25, 35, 38, 58, 60) and the vertical transmission of microbes from the mother (75–77). However, we detected microorganisms present in fetal samples but absent in the maternal intestine. Several studies suggested that the fetal microbiome may be attributed to multiple potential maternal sources, including oral, endometrial, and urogenital microbiotas (78–82). Such a diverse range of microbiota sources could account for the difference in microorganisms observed between the maternal intestine and fetal samples. Together, our analysis provides significant insights into the microbiota and immune development of rhesus monkey fetuses.

However, the presence of fetal microbiota remains controversial, with the long-held view positing the fetus develops in a sterile intrauterine environment until birth (83). Moreover, a recent study by Theis et al. isolated *Cutibacterium acnes* from one colony out of 96 fetal and placental samples of RMs, and with 16S rRNA sequencing, they found the bacterium in the fetal samples, and particularly in the maternal decidua with relative high abundance. However, given that this bacterium was also detected in nearly half of the background technical controls and is commonly present on human skin (84) there is a possibility of contamination (57). This study obtained a RM fetus in a sterile manner by cesarean section without contamination from the birth canal, and we applied stringent experimental conditions and rigorous control settings to avoid contamination. We also detected the *Cutibacterium acnes* with relatively high abundance in the fetal samples. Consistent with our result, *Cutibacterium acnes* was reported to be a significant portion of bacterial isolates from human placenta and amniotic fluid in normal term pregnancies (85). Unfortunately, due to the limited sample size, whether *Cutibacterium acnes* exists in the intrauterine environment of RM or not, still needs further verification with advanced analysis methods and more fetal samples in future. Different to the case of *Cutibacterium acnes,* other bacteria such as *Acinetobacter* and *Ottowia,* identified in our fetal samples, were also detected in uterine wall samples of RMs but were seldom found in controls by Theis et al’s study (57). These bacteria have also been detected from the human endometrium (86–91). Additionally, *Ralstonia insidiosa*, found in high abundance in our fetal samples, was a resident bacterium at the maternal-fetal interface (40). This bacterium was detected in the human placental basal plate and villus through 16S rRNA sequencing and validated by quantitative real-time PCR and fluorescent in situ hybridization (39, 40). Parnell et al. further demonstrated that *Ralstonia insidiosa* can enter the placenta via the intrauterine route in a pregnant mouse model, suggesting a mechanism for its presence at the maternal-fetal interface (40). This evidence supports their relevance to the uterine environment and reinforces our hypothesis about their presence in the RM uterine environment.

Recent studies have highlighted the functional involvement of archaea, single-celled prokaryotes, in health and disease (92, 93). Our results indicated that the most abundant archaea at the phylum level, Euryarchaeota, was shared by maternal and fetal samples. Euryarchaeota contains the greatest number and diversity of archaea (94), including methanogens. The methanogens were also the dominant archaeal component in fetal samples in our study. At present, however, the functional and pathogenic impacts of archaea in animals remain poorly understood (95, 96). Additionally, the viral profiles in both maternal and fetal samples were similar, indicating the possibility of vertical transmission of viruses from the mother to the fetus within the uterus.

### 4.2 Microbes in placenta and umbilical cord

Identification of microbes in the placenta and umbilical cord is crucial, as the placenta serves as the primary barrier between the fetus and mother, and the umbilical cord supplies essential nutrients for fetal development and survival (97, 98). However, the mechanism by which microbes traverse the placental barrier and reach the fetus remains unresolved, despite the potential impact of microbes on fetal health (99, 100). Studies on the presence of microorganisms in the placenta and umbilical cord are pivotal for developing effective strategies to prevent fetal microbial infections.

In the present study, diverse microbes were identified in the placenta, umbilical cord, and two fetal organs, while the umbilical cord exhibited a lower microbial DNA content relative to other samples. The microbial profiles of the placenta and umbilical cord more closely resembled those of the fetal spleen and fetal GI than the maternal intestine. The placenta, umbilical cord, and fetal organs shared various dominant microbes and showed similar microbial diversity and composition, strongly suggesting that placental microbes may be the primary contributors to the fetal microbiome (25, 55, 74). Proteobacteria was the most abundant bacteria in the placenta, consistent with findings from human studies (85, 101). The umbilical cord microbiota differed somewhat from that of other fetal samples, possibly due to its transportation role, relatively low microbial DNA content, or increased host contamination. *Staphylococcus* was notably abundant in the umbilical cord samples. This bacterium has been detected in fetal and placental samples from Japanese macaques (55), as well as in human amniotic fluid, placental, and cord blood samples (85, 102). While prevalent in human placenta and various fetal organs through clinical culturing (35), some sequence-based studies suggest their presence may be due to background contamination (103).

### 4.3 Microbial differences between adult RMs and fetus

Physiological and nutritional changes from the fetal stage to adulthood induce persistent changes in the microbial composition and enrichment of genes associated with carbohydrate metabolism (104, 105). In contrast to fetal nutrition, which relies on glucose provided by the mother (106), adult RMs encounter a much more complex environment and diet. Our results showed pronounced differences in microbial compositions and functions between adult RMs and fetuses. Regarding microbial composition, the abundant microbial taxa differed between adult and fetal samples, with adults exhibiting significantly higher microbial richness. In addition, the different bacteria possess carbohydrates targeting distinct polysaccharides for degradation and facilitate their colonization in the gut through the evolution of carbohydrates (107, 108). Consequently, adult RMs may exhibit a more pronounced carbohydrate metabolism than fetuses, as reflected by our results showing the greater diversity and abundance of CAZyme genes detected in the adult microbiome. The dominant CAZymes in the adult and fetal microbiomes were different. In the adult microbiome, GHs were the predominant CAZymes, while in fetuses, both GHs and GTs were prevalent. These enzyme families are primarily responsible for carbohydrate degradation and synthesis, respectively (109, 110). GH family genes were significantly enriched in adult microbiome than fetal microbiome, especially GH1 and GH43, which are associated with hemicellulose and cellulose degradation (109–111) and support the dietary demands of adult RMs to metabolize fiber-intensive foods (112, 113). In both the adult and fetal microbiomes, GH13 was the dominant GH family. Notably, GH13, a significant family of glycoside hydrolases, is involved in the degradation of glycogen, oligosaccharides, polysaccharides, and starch (114, 115).

Maternal antibiotic treatment during pregnancy has long-lasting effects on the fetal microbiota and health (116–118). Maternal ARGs can be transmitted to the fetus and influence the fetal microbiome and resistome profiles to induce resistance (119–121). As the fetal microbiota is a critical determinant of early immunity development, overall health, and antibiotic treatment efficacy (35), a comprehensive understanding of fetal ARGs is crucial. In this study, even without direct antibiotic exposure to the fetus, we identified four ARGs in the fetal samples that aligned with those in maternal and adult RMs, suggesting a vertical transmission of ARG-containing microbes during pregnancy. Subsequent examinations of bacteria harboring ARGs, and the impact of specific antibiotics may aid in the development of preventive and treatment strategies for healthy pregnancy outcomes (122–124).

### 4.4 Microbial differences between human and RM fetuses

The microbial components of NHPs tend to resemble those of humans more closely than those of other animals (125, 126). Our analysis revealed a convergence in certain dominant microbiota between human and RM fetuses. Specifically, Proteobacteria was the most abundant phylum in both human fetal intestines and RM fetal organs. Other research has highlighted the dominance of Proteobacteria in the human neonatal gut during the first week of life (127), which facilitates colonization by strict anaerobes and underscores the susceptibility of bacterial communities at this stage (128, 129). Proteobacteria constitutes a major community in the human placenta and umbilical cord blood (37, 130), with overlaps noted between placental communities and those in amniotic fluid and meconium (131, 132). Additionally, the intestinal abundance of Proteobacteria in both macaques and humans significantly increases during pregnancy (133, 134). Based on published studies and the high abundance of Proteobacteria observed in maternal RM in our study, it is reasonable to speculate that, similar to humans (135), Proteobacteria in RM fetuses may also be transmitted from the mother *in utero*. Of note, Firmicutes and Bacteroidetes are reported to be the primary phyla in both adult humans and RMs (136–140), suggesting that, during the transition from fetus to adulthood in both species, Firmicutes and Bacteroidetes displace Proteobacteria as the prevailing bacteria. Our results also revealed that methanogens were the dominant archaea in the fetuses of both human and RM. These observations underscore the similarity in microbial communities between human and RM fetuses, suggesting macaques as good models to study human microbial composition and transmission.

However, disparities in the microbiome between adult humans and RMs were evident, likely stemming from significant differences in genetic, physiological, dietary, environmental factors, among others (141–143). In our microbial analysis, significant differences were observed in the overall compositions, potential functions, and abundance of specific microbes between the human and RM fetuses. Given the microbial differences between RMs and humans, caution is necessary when extrapolating the findings of this study to human biology.

In conclusion, our research reported a detailed case analysis of the microbial composition and function within an RM fetus, meticulously obtained via cesarean section to avoid contamination from the birth canal. By preparing DNA libraries from environmental and blank controls, we confirmed low environmental contamination in our samples, reinforcing the validity of our findings. We observed a diverse array of microorganisms in the womb and fetus. As crucial connections between mother and fetus, the placenta and umbilical cord exhibit a microbial composition that more closely resembles that of the fetus rather than the mother. These observations suggest a potential maternal-fetal microbial transfer, possibly through these structures, as supported by substantial microbial sharing and clustered microbial profiles between the mother and fetus compared to other adult RMs. Importantly, the discovery of seven ARGs in the RM fetus further suggests potential vertical transmission, highlighting the possible critical role these microbes may play in fetal health and immune system development. In terms of the unique fetal microbial profile, our findings reveal substantial differences in microbial composition and functional pathways between the fetus and adults. Notably, the fetus exhibited a less developed capacity for carbohydrate metabolism, characterized by fewer diverse genes and less complex pathways compared to adults. Our findings not only emphasize maternal-fetal microbial transfer but also help in formulating strategies to tackle microbial-induced challenges in fetal health and development. However, it is noteworthy that, obtaining placental and fetal samples from NHPs is known to be difficult due to ethical and welfare considerations. Future research, enriched with more microbiome data from macaque fetal samples and innovative methodologies, will benefit to elucidate the impact of fetal microbiota on the developing immune system, thereby advancing our understanding of these complex biological processes.

## Declarations

## Data availability

The raw sequence data reported in this paper have been deposited in the Genome Sequence Archive in National Genomics Data Center (GSA: CRA014938 and CRA014939) that are accessible at https://ngdc.cncb.ac.cn/gsa.

## Ethics approval and consent to participate

This study was approved by the Ethics Committee of the College of Life Sciences, Sichuan University, China (SCU230810001). All sample collection and utility protocols fully complied with the guidelines of the Management Committee of Experimental Animals of Sichuan Province, China (SYXK-Sichuan, 2019-192).

## Consent for publication

Not Applicable.

## Authors’ contributions

QD: Writing – original draft, Writing – review & editing, Formal Analysis, Visualization; XL: Writing – review & editing, Methodology, Formal Analysis; RSZ: Writing – review & editing, Methodology; GH: Writing – review & editing, Data curation; Resources; QHL: Writing – review & editing, Data curation; Resources; WM: Writing – review & editing, Visualization; YH: Writing – review & editing, Visualization; ZXF: Writing – review & editing, Project administration; JL: Supervision, Writing – review & editing, Funding acquisition.

## Acknowledgements

We thank Novogene Bioinformatics Technology Co., Ltd. (Beijing, China) for metagenomic sequencing and give special thanks to Sichuan Green-House Biotech Co., Ltd. for the provision of samples.

## References

1. Halkias J., Rackaityte E, Hillman SL, Aran D, Mendoza VF, Marshall LR, et al. Cd161 Contributes to Prenatal Immune Suppression of Ifn-Γ–Producing Plzf+ T Cells. The Journal of Clinical Investigation (2019) 129(9):3562–77.

2. Le BL, Sper R, Nielsen SC, Pineda S, Nguyen Q-H, Lee J-Y, et al. Maternal and Infant Immune Repertoire Sequencing Analysis Identifies Distinct Ig and Tcr Development in Term and Preterm Infants. The Journal of Immunology (2021) 207(10):2445–55.

3. McGovern N, Shin A, Low G, Low D, Duan K, Yao LJ, et al. Human Fetal Dendritic Cells Promote Prenatal T-Cell Immune Suppression through Arginase-2. Nature (2017) 546(7660):662–6.

4. Papadopoulou M, Tieppo P, McGovern N, Gosselin F, Chan JK, Goetgeluk G, et al. Tcr Sequencing Reveals the Distinct Development of Fetal and Adult Human Vγ9vδ2 T Cells. The Journal of Immunology (2019) 203(6):1468–79.

5. Park J-E, Botting RA, Domínguez Conde C, Popescu D-M, Lavaert M, Kunz DJ, et al. A Cell Atlas of Human Thymic Development Defines T Cell Repertoire Formation. Science (2020) 367(6480):eaay3224.

6. Rackaityte E, Halkias J. Mechanisms of Fetal T Cell Tolerance and Immune Regulation. Frontiers in immunology (2020) 11:588.

7. Romero R, Gomez R, Ghezzi F, Yoon BH, Mazor M, Edwin SS, et al. A Fetal Systemic Inflammatory Response Is Followed by the Spontaneous Onset of Preterm Parturition. American journal of obstetrics and gynecology (1998) 179(1):186–93.

8. Zhivaki D, Lo-Man R, editors. In Utero Development of Memory T Cells. Seminars in Immunopathology; 2017: Springer.

9. Rechavi E, Lev A, Lee YN, Simon AJ, Yinon Y, Lipitz S, et al. Timely and Spatially Regulated Maturation of B and T Cell Repertoire During Human Fetal Development. Science translational medicine (2015) 7(276):276ra25–ra25.

10. Leavy O. Fetal Immune Repertoire. Nature Reviews Immunology (2015) 15(4):201–. doi: 10.1038/nri3841.

11. Para R, Romero R, Miller D, Galaz J, Done B, Peyvandipour A, et al. The Distinct Immune Nature of the Fetal Inflammatory Response Syndrome Type I and Type Ii. ImmunoHorizons (2021) 5(9):735–51.

12. Gotsch F, Romero R, Kusanovic JP, Mazaki-Tovi S, Pineles BL, Erez O, et al. The Fetal Inflammatory Response Syndrome. Clinical obstetrics and gynecology (2007) 50(3):652–83.

13. Li N, van Unen V, Abdelaal T, Guo N, Kasatskaya SA, Ladell K, et al. Memory Cd4+ T Cells Are Generated in the Human Fetal Intestine. Nature immunology (2019) 20(3):301–12.

14. Nikiforou M, Jacobs EM, Kemp MW, Hornef MW, Payne MS, Saito M, et al. Intra-Amniotic Candida Albicans Infection Induces Mucosal Injury and Inflammation in the Ovine Fetal Intestine. Scientific reports (2016) 6(1):1–9.

15. Schreurs RR, Baumdick ME, Sagebiel AF, Kaufmann M, Mokry M, Klarenbeek PL, et al. Human Fetal Tnf-Α-Cytokine-Producing Cd4+ Effector Memory T Cells Promote Intestinal Development and Mediate Inflammation Early in Life. Immunity (2019) 50(2):462–76. e8.

16. Stras SF, Werner L, Toothaker JM, Olaloye OO, Oldham AL, McCourt CC, et al. Maturation of the Human Intestinal Immune System Occurs Early in Fetal Development. Developmental cell (2019) 51(3):357–73. e5.

17. Wolfs TG, Kramer BW, Thuijls G, Kemp MW, Saito M, Willems MG, et al. Chorioamnionitis-Induced Fetal Gut Injury Is Mediated by Direct Gut Exposure of Inflammatory Mediators or by Lung Inflammation. American Journal of Physiology-Gastrointestinal and Liver Physiology (2014) 306(5):G382–G93.

18. Robbins JR, Bakardjiev AI. Pathogens and the Placental Fortress. Current Opinion in Microbiology (2012) 15(1):36–43. doi: 10.1016/j.mib.2011.11.006.

19. Delorme-Axford E, Donker RB, Mouillet J-F, Chu T, Bayer A, Ouyang Y, et al. Human Placental Trophoblasts Confer Viral Resistance to Recipient Cells. Proceedings of the National Academy of Sciences (2013) 110(29):12048–53.

20. Bayer A, Delorme-Axford E, Sleigher C, Frey TK, Trobaugh DW, Klimstra WB, et al. Human Trophoblasts Confer Resistance to Viruses Implicated in Perinatal Infection. American journal of obstetrics and gynecology (2015) 212(1):71. e1-. e8.

21. Liu Y, Gao S, Zhao Y, Wang H, Pan Q, Shao Q. Decidual Natural Killer Cells: A Good Nanny at the Maternal-Fetal Interface During Early Pregnancy. Frontiers in Immunology (2021):1684.

22. PrabhuDas M, Bonney E, Caron K, Dey S, Erlebacher A, Fazleabas A, et al. Immune Mechanisms at the Maternal-Fetal Interface: Perspectives and Challenges. Nature immunology (2015) 16(4):328–34.

23. Than NG, Hahn S, Rossi SW, Szekeres-Bartho J. Fetal-Maternal Immune Interactions in Pregnancy. Frontiers Media SA (2019). p. 2729.

24. Al Alam D, Danopoulos S, Grubbs B, Ali NAtBM, MacAogain M, Chotirmall SH, et al. Human Fetal Lungs Harbor a Microbiome Signature. American journal of respiratory and critical care medicine (2020) 201(8):1002–6.

25. Younge N, McCann JR, Ballard J, Plunkett C, Akhtar S, Araújo-Pérez F, et al. Fetal Exposure to the Maternal Microbiota in Humans and Mice. JCI insight (2019) 4(19).

26. Willyard C. Baby’s First Bacteria. Nature (2018) 553:264–6.

27. Seferovic MD, Pace RM, Carroll M, Belfort B, Major AM, Chu DM, et al. Visualization of Microbes by 16s in Situ Hybridization in Term and Preterm Placentas without Intraamniotic Infection. American Journal of Obstetrics and Gynecology (2019) 221(2):146. e1-. e23.

28. Rackaityte E, Halkias J, Fukui E, Mendoza V, Hayzelden C, Crawford E, et al. Viable Bacterial Colonization Is Highly Limited in the Human Intestine in Utero. Nature medicine (2020) 26(4):599–607.

29. Perez-Muñoz ME, Arrieta M-C, Ramer-Tait AE, Walter J. A Critical Assessment of the “Sterile Womb” and “in Utero Colonization” Hypotheses: Implications for Research on the Pioneer Infant Microbiome. microbiome (2017) 5(1):1–19.

30. Kundu P, Blacher E, Elinav E, Pettersson S. Our Gut Microbiome: The Evolving Inner Self. Cell (2017) 171(7):1481–93.

31. Heijtz RD, editor. Fetal, Neonatal, and Infant Microbiome: Perturbations and Subsequent Effects on Brain Development and Behavior. Seminars in Fetal and Neonatal Medicine; 2016: Elsevier.

32. Hussain T, Murtaza G, Kalhoro DH, Kalhoro MS, Yin Y, Chughtai MI, et al. Understanding the Immune System in Fetal Protection and Maternal Infections During Pregnancy. Journal of Immunology Research (2022) 2022.

33. Jain N. The Early Life Education of the Immune System: Moms, Microbes and (Missed) Opportunities. Gut Microbes (2020) 12(1):1824564.

34. McCauley KE, Rackaityte E, LaMere B, Fadrosh DW, Fujimura KE, Panzer AR, et al. Heritable Vaginal Bacteria Influence Immune Tolerance and Relate to Early-Life Markers of Allergic Sensitization in Infancy. Cell Reports Medicine (2022) 3(8):100713.

35. Mishra A, Lai GC, Yao LJ, Aung TT, Shental N, Rotter-Maskowitz A, et al. Microbial Exposure During Early Human Development Primes Fetal Immune Cells. Cell (2021) 184(13):3394–409. e20.

36. Vidal Jr MS, Menon R. In Utero Priming of Fetal Immune Activation: Myths and Mechanisms. Journal of Reproductive Immunology (2023):103922.

37. Aagaard K, Ma J, Antony KM, Ganu R, Petrosino J, Versalovic J. The Placenta Harbors a Unique Microbiome. Science translational medicine (2014) 6(237):237ra65–ra65.

38. Leon LJ, Doyle R, Diez-Benavente E, Clark TG, Klein N, Stanier P, et al. Enrichment of Clinically Relevant Organisms in Spontaneous Preterm-Delivered Placentas and Reagent Contamination across All Clinical Groups in a Large Pregnancy Cohort in the United Kingdom. Applied and environmental microbiology (2018) 84(14):e00483–18.

39. Parnell LA, Briggs CM, Cao B, Delannoy-Bruno O, Schrieffer AE, Mysorekar IU. Microbial Communities in Placentas from Term Normal Pregnancy Exhibit Spatially Variable Profiles. Scientific reports (2017) 7(1):11200.

40. Parnell LA, Willsey GG, Joshi CS, Yin Y, Wargo MJ, Mysorekar IU. Functional Characterization of Ralstonia Insidiosa, a Bona Fide Resident at the Maternal-Fetal Interface. bioRxiv (2019):721977.

41. Egli G, Newton M. The Transport of Carbon Particles in the Human Female Reproductive Tract. Fertility and sterility (1961) 12(2):151–5.

42. Seong HS, Lee SE, Kang JH, Romero R, Yoon BH. The Frequency of Microbial Invasion of the Amniotic Cavity and Histologic Chorioamnionitis in Women at Term with Intact Membranes in the Presence or Absence of Labor. American journal of obstetrics and gynecology (2008) 199(4):375. e1-. e5.

43. Romero R, Nores J, Mazor M, Sepulveda W, Oyarzun E, Parra M, et al. Microbial Invasion of the Amniotic Cavity During Term Labor. Prevalence and Clinical Significance. The Journal of reproductive medicine (1993) 38(7):543–8.

44. Grigsby PL, editor. Animal Models to Study Placental Development and Function Throughout Normal and Dysfunctional Human Pregnancy. Seminars in reproductive medicine; 2016: Thieme Medical Publishers.

45. Pronovost GN, Yu KB, Coley-O’Rourke EJ, Telang SS, Chen AS, Vuong HE, et al. The Maternal Microbiome Promotes Placental Development in Mice. Science Advances (2023) 9(40):eadk1887.

46. Phillips KA, Bales KL, Capitanio JP, Conley A, Czoty PW, ‘t Hart BA, et al. Why Primate Models Matter. American journal of primatology (2014) 76(9):801–27.

47. Carlsson HE, Schapiro SJ, Farah I, Hau J. Use of Primates in Research: A Global Overview. American Journal of Primatology: Official Journal of the American Society of Primatologists (2004) 63(4):225–37.

48. Carter AM. Animal Models of Human Placentation–a Review. Placenta (2007) 28:S41–S7.

49. Buse E, Häeger J-D, Svensson-Arvelund J, Markert UR, Faas MM, Ernerudh J, et al. The Placenta in Toxicology. Part I: Animal Models in Toxicology: Placental Morphology and Tolerance Molecules in the Cynomolgus Monkey (Macaca Fascicularis). Toxicologic Pathology (2014) 42(2):314–26.

50. Estes JD, Wong SW, Brenchley JM. Nonhuman Primate Models of Human Viral Infections. Nature reviews Immunology (2018) 18(6):390–404.

51. Daadi MM, Barberi T, Shi Q, Lanford RE. Nonhuman Primate Models in Translational Regenerative Medicine. Stem Cells and Development (2014) 23(S1):83–7.

52. Newman C, Friedrich TC, O’Connor DH. Macaque Monkeys in Zika Virus Research: 1947– Present. Current opinion in virology (2017) 25:34–40.

53. Prescott MJ. Ethics of Primate Use. Advances in science and research (2010) 5(1):11–22.

54. Vermeire T, Epstein M, Badin R, Flecknell P, Hoet P, Hudson-Shore M, et al. Final Opinion on the Need for Non-Human Primates in Biomedical Research, Production and Testing of Products and Devices (Update 2017). Sci Comm Heal Environ Emerg Risks (2017).

55. Chu D, Prince A, Ma J, Baquero K, Blundell P, Takahashi D, et al. 114: Evidence of Fetal Microbiota and Its Maternal Origins in a Non-Human Primate Model. American Journal of Obstetrics & Gynecology (2017) 216(1):S80.

56. Prince A, Chu D, Meyer K, Ma J, Baquero K, Blundell P, et al. 23: The Fetal Microbiome Is Altered in Association with Maternal Diet During Gestation. American Journal of Obstetrics & Gynecology (2017) 216(1):S17.

57. Theis KR, Romero R, Winters AD, Jobe AH, Gomez-Lopez N. Lack of Evidence for Microbiota in the Placental and Fetal Tissues of Rhesus Macaques. Msphere (2020) 5(3):e00210–20.

58. Chu D, Prince A, Ma J, Pace R, Takahashi D, Friedman J, et al. 115: Contribution of the Fetal Microbiome to the Taxonomic Diversity and Functionality of the Postnatal Gut Microbiome in a Non-Human Primate (Nhp) Model. American Journal of Obstetrics & Gynecology (2018) 218(1):S82–S3.

59. Prince A, Ma J, Hu M, Chu D, Miller L, Jobe A, et al. 843: Chorioamnionitis Induced by Intra-Amniotic Injection of Il-1, Lps, or Ureaplasma Parvum Is Associated with an Altered Microbiome in a Primate Model of Inflammatory Preterm Birth. American Journal of Obstetrics & Gynecology (2018) 218(1):S503.

60. Prince AL, Ma J, Hu M, Chu DM, Miller L, Jobe A, et al. 521: Intra-Amniotic Injection Alters the Intrauterine Microbiome in a Primate Model of Inflammatory Preterm Birth. American Journal of Obstetrics & Gynecology (2019) 220(1):S349.

61. Chen S, Zhou Y, Chen Y, Gu J. Fastp: An Ultra-Fast All-in-One Fastq Preprocessor. Bioinformatics (2018) 34(17):i884–i90.

62. Langmead B, Salzberg SL. Fast Gapped-Read Alignment with Bowtie 2. Nature methods (2012) 9(4):357–9.

63. Li D, Liu C-M, Luo R, Sadakane K, Lam T-W. Megahit: An Ultra-Fast Single-Node Solution for Large and Complex Metagenomics Assembly Via Succinct De Bruijn Graph. Bioinformatics (2015) 31(10):1674–6.

64. Hyatt D, Chen G-L, LoCascio PF, Land ML, Larimer FW, Hauser LJ. Prodigal: Prokaryotic Gene Recognition and Translation Initiation Site Identification. BMC bioinformatics (2010) 11:1–11.

65. Fu L, Niu B, Zhu Z, Wu S, Li W. Cd-Hit: Accelerated for Clustering the Next-Generation Sequencing Data. Bioinformatics (2012) 28(23):3150–2.

66. Patro R, Duggal G, Love MI, Irizarry RA, Kingsford C. Salmon Provides Fast and Bias-Aware Quantification of Transcript Expression. Nature methods (2017) 14(4):417–9.

67. Buchfink B, Xie C, Huson DH. Fast and Sensitive Protein Alignment Using Diamond. Nature methods (2015) 12(1):59–60.

68. Lombard V, Golaconda Ramulu H, Drula E, Coutinho PM, Henrissat B. The Carbohydrate-Active Enzymes Database (Cazy) in 2013. Nucleic acids research (2014) 42(D1):D490–D5.

69. Alcock BP, Raphenya AR, Lau TT, Tsang KK, Bouchard M, Edalatmand A, et al. Card 2020: Antibiotic Resistome Surveillance with the Comprehensive Antibiotic Resistance Database. Nucleic acids research (2020) 48(D1):D517–D25.

70. Franzosa EA, McIver LJ, Rahnavard G, Thompson LR, Schirmer M, Weingart G, et al. Species-Level Functional Profiling of Metagenomes and Metatranscriptomes. Nature methods (2018) 15(11):962–8.

71. Suzek BE, Huang H, McGarvey P, Mazumder R, Wu CH. Uniref: Comprehensive and Non-Redundant Uniprot Reference Clusters. Bioinformatics (2007) 23(10):1282–8.

72. Wood DE, Salzberg SL. Kraken: Ultrafast Metagenomic Sequence Classification Using Exact Alignments. Genome biology (2014) 15(3):1–12.

73. R Core Team R. R: A Language and Environment for Statistical Computing. (2013).

74. Cao B, Stout MJ, Lee I, Mysorekar IU. Placental Microbiome and Its Role in Preterm Birth. Neoreviews (2014) 15(12):e537–e45.

75. Hirsch AJ, Roberts VH, Grigsby PL, Haese N, Schabel MC, Wang X, et al. Zika Virus Infection in Pregnant Rhesus Macaques Causes Placental Dysfunction and Immunopathology. Nature communications (2018) 9(1):263.

76. Aronsson F, Lannebo C, Paucar M, Brask J, Kristensson K, Karlsson H. Persistence of Viral Rna in the Brain of Offspring to Mice Infected with Influenza a/Wsn/33 Virus During Pregnancy. Journal of neurovirology (2002) 8(4):353–7.

77. Wolfe B, Kerr AR, Mejia A, Simmons HA, Czuprynski CJ, Golos TG. Sequelae of Fetal Infection in a Non-Human Primate Model of Listeriosis. Frontiers in Microbiology (2019) 10:2021.

78. Walker RW, Clemente JC, Peter I, Loos RJF. The Prenatal Gut Microbiome: Are We Colonized with Bacteria in Utero? Pediatr Obes (2017) 12 Suppl 1(Suppl 1):3–17. Epub 2017/04/28. doi: 10.1111/ijpo.12217.

79. Yu K, Rodriguez M, Paul Z, Gordon E, Gu T, Rice K, et al. Transfer of Oral Bacteria to the Fetus During Late Gestation. Scientific Reports (2021) 11(1):708. doi: 10.1038/s41598-020-80653-y.

80. Keelan JA, Payne MS. Vaginal Microbiota During Pregnancy: Pathways of Risk of Preterm Delivery in the Absence of Intrauterine Infection? Proceedings of the National Academy of Sciences (2015) 112(47):E6414–E.

81. Goldenberg RL, Hauth JC, Andrews WW. Intrauterine Infection and Preterm Delivery. New England journal of medicine (2000) 342(20):1500–7.

82. Stout MJ, Conlon B, Landeau M, Lee I, Bower C, Zhao Q, et al. Identification of Intracellular Bacteria in the Basal Plate of the Human Placenta in Term and Preterm Gestations. American journal of obstetrics and gynecology (2013) 208(3):226. e1-. e7.

83. Walter J, Hornef MW. A Philosophical Perspective on the Prenatal in Utero Microbiome Debate. Microbiome (2021) 9(1):5.

84. Omer H, McDowell A, Alexeyev OA. Understanding the Role of Propionibacterium Acnes in Acne Vulgaris: The Critical Importance of Skin Sampling Methodologies. Clinics in dermatology (2017) 35(2):118–29.

85. Collado MC, Rautava S, Aakko J, Isolauri E, Salminen S. Human Gut Colonisation May Be Initiated in Utero by Distinct Microbial Communities in the Placenta and Amniotic Fluid. Scientific reports (2016) 6(1):1–13.

86. Devi CA, Ranjani A, Dhanasekaran D, Thajuddin N, Ramanidevi T. Surveillance of Multidrug Resistant Bacteria Pathogens from Female Infertility Cases. African Journal of Biotechnology (2013) 12(26).

87. Moreno I, Codoñer FM, Vilella F, Valbuena D, Martinez-Blanch JF, Jimenez-Almazán J, et al. Evidence That the Endometrial Microbiota Has an Effect on Implantation Success or Failure. American journal of obstetrics and gynecology (2016) 215(6):684–703.

88. Winters AD, Romero R, Gervasi MT, Gomez-Lopez N, Tran MR, Garcia-Flores V, et al. Does the Endometrial Cavity Have a Molecular Microbial Signature? Scientific reports (2019) 9(1):9905.

89. Leoni C, Ceci O, Manzari C, Fosso B, Volpicella M, Ferrari A, et al. Human Endometrial Microbiota at Term of Normal Pregnancies. Genes (2019) 10(12):971.

90. Chen C, Song X, Wei W, Zhong H, Dai J, Lan Z, et al. The Microbiota Continuum Along the Female Reproductive Tract and Its Relation to Uterine-Related Diseases. Nature communications (2017) 8(1):875.

91. Miles SM, Hardy BL, Merrell DS. Investigation of the Microbiota of the Reproductive Tract in Women Undergoing a Total Hysterectomy and Bilateral Salpingo-Oopherectomy. Fertility and sterility (2017) 107(3):813–20. e1.

92. Chehoud C, Albenberg LG, Judge C, Hoffmann C, Grunberg S, Bittinger K, et al. Fungal Signature in the Gut Microbiota of Pediatric Patients with Inflammatory Bowel Disease. Inflammatory bowel diseases (2015) 21(8):1948–56.

93. Lewis JD, Chen EZ, Baldassano RN, Otley AR, Griffiths AM, Lee D, et al. Inflammation, Antibiotics, and Diet as Environmental Stressors of the Gut Microbiome in Pediatric Crohn’s Disease. Cell host & microbe (2015) 18(4):489–500.

94. Baker BJ, De Anda V, Seitz KW, Dombrowski N, Santoro AE, Lloyd KG. Diversity, Ecology and Evolution of Archaea. Nature microbiology (2020) 5(7):887–900.

95. Kumondorova A, Serkan İ. Archaea and Their Potential Pathogenicity in Human and Animal Diseases. Journal of Istanbul Veterinary Sciences (2019) 3(3):79–84.

96. Cavicchioli R, Curmi PM, Saunders N, Thomas T. Pathogenic Archaea: Do They Exist? Bioessays (2003) 25(11):1119–28.

97. Sapunar D, Arey LB, Rogers K. Prenatal Development (2023) [cited 2022 October 18]. Available from: https://www.britannica.com/science/prenatal-development.

98. Macpherson AJ, de Agüero MG, Ganal-Vonarburg SC. How Nutrition and the Maternal Microbiota Shape the Neonatal Immune System. Nature Reviews Immunology (2017) 17(8):508–17.

99. Megli CJ, Coyne CB. Infections at the Maternal–Fetal Interface: An Overview of Pathogenesis and Defence. Nature Reviews Microbiology (2022) 20(2):67–82. doi: 10.1038/s41579-021-00610-y.

100. Arora N, Sadovsky Y, Dermody TS, Coyne CB. Microbial Vertical Transmission During Human Pregnancy. Cell Host Microbe (2017) 21(5):561–7. Epub 2017/05/12. doi: 10.1016/j.chom.2017.04.007.

101. Nuriel-Ohayon M, Neuman H, Koren O. Microbial Changes During Pregnancy, Birth, and Infancy. Frontiers in microbiology (2016):1031.

102. Jiménez E, Fernández L, Marín ML, Martín R, Odriozola JM, Nueno-Palop C, et al. Isolation of Commensal Bacteria from Umbilical Cord Blood of Healthy Neonates Born by Cesarean Section. Curr Microbiol (2005) 51(4):270–4. Epub 2005/09/28. doi: 10.1007/s00284-005-0020-3.

103. Glassing A, Dowd SE, Galandiuk S, Davis B, Chiodini RJ. Inherent Bacterial DNA Contamination of Extraction and Sequencing Reagents May Affect Interpretation of Microbiota in Low Bacterial Biomass Samples. Gut pathogens (2016) 8:1–12.

104. Koenig JE, Spor A, Scalfone N, Fricker AD, Stombaugh J, Knight R, et al. Succession of Microbial Consortia in the Developing Infant Gut Microbiome. Proceedings of the National Academy of Sciences (2011) 108(supplement_1):4578–85.

105. Wardman JF, Bains RK, Rahfeld P, Withers SG. Carbohydrate-Active Enzymes (Cazymes) in the Gut Microbiome. Nature Reviews Microbiology (2022) 20(9):542–56.

106. Kalhan S, Parimi P. Gluconeogenesis in the Fetus and Neonate. Semin Perinatol (2000) 24(2):94–106. Epub 2000/05/11. doi: 10.1053/sp.2000.6360.

107. Martens EC, Kelly AG, Tauzin AS, Brumer H. The Devil Lies in the Details: How Variations in Polysaccharide Fine-Structure Impact the Physiology and Evolution of Gut Microbes. Journal of molecular biology (2014) 426(23):3851–65.

108. Kaoutari AE, Armougom F, Gordon JI, Raoult D, Henrissat B. The Abundance and Variety of Carbohydrate-Active Enzymes in the Human Gut Microbiota. Nature Reviews Microbiology (2013) 11(7):497–504.

109. Comtet-Marre S, Chaucheyras-Durand F, Bouzid O, Mosoni P, Bayat AR, Peyret P, et al. Fibrochip, a Functional DNA Microarray to Monitor Cellulolytic and Hemicellulolytic Activities of Rumen Microbiota. Frontiers in microbiology (2018) 9:215.

110. Kuntothom T, Luang S, Harvey AJ, Fincher GB, Opassiri R, Hrmova M, et al. Rice Family Gh1 Glycoside Hydrolases with Β-D-Glucosidase and Β-D-Mannosidase Activities. Archives of Biochemistry and Biophysics (2009) 491(1-2):85–95.

111. Heins RA, Cheng X, Nath S, Deng K, Bowen BP, Chivian DC, et al. Phylogenomically Guided Identification of Industrially Relevant Gh1 Β-Glucosidases through DNA Synthesis and Nanostructure-Initiator Mass Spectrometry. ACS chemical biology (2014) 9(9):2082–91.

112. Sun B, Xia Y, Garber PA, Amato KR, Gomez A, Xu X, et al. Captivity Is Associated with Gut Mycobiome Composition in Tibetan Macaques (Macaca Thibetana). Frontiers in Microbiology (2021) 12:665853.

113. Wang Y, Yang X, Zhang M, Pan H. Comparative Analysis of Gut Microbiota between Wild and Captive Golden Snub-Nosed Monkeys. Animals (2023) 13(10):1625.

114. Kuriki T, Imanaka T. The Concept of the Α-Amylase Family: Structural Similarity and Common Catalytic Mechanism. Journal of bioscience and bioengineering (1999) 87(5):557–65.

115. Stam MR, Danchin EG, Rancurel C, Coutinho PM, Henrissat B. Dividing the Large Glycoside Hydrolase Family 13 into Subfamilies: Towards Improved Functional Annotations of Α-Amylase-Related Proteins. Protein Engineering, Design and Selection (2006) 19(12):555–62.

116. Nahum GG, Uhl K, Kennedy DL. Antibiotic Use in Pregnancy and Lactation: What Is and Is Not Known About Teratogenic and Toxic Risks. Obstetrics & Gynecology (2006) 107(5):1120–38.

117. Prescott S, Dreisbach C, Baumgartel K, Koerner R, Gyamfi A, Canellas M, et al. Impact of Intrapartum Antibiotic Prophylaxis on Offspring Microbiota. Frontiers in Pediatrics (2021) 9:754013.

118. Chen C-M, Chou H-C, Yang Y-CS. Maternal Antibiotic Treatment Disrupts the Intestinal Microbiota and Intestinal Development in Neonatal Mice. Frontiers in Microbiology (2021) 12:684233.

119. Patangia DV, Ryan CA, Dempsey E, Stanton C, Ross RP. Vertical Transfer of Antibiotics and Antibiotic Resistant Strains across the Mother/Baby Axis. Trends in Microbiology (2022) 30(1):47–56.

120. Bi Y, Tu Y, Zhang N, Wang S, Zhang F, Suen G, et al. Multiomics Analysis Reveals the Presence of a Microbiome in the Gut of Fetal Lambs. Gut (2021) 70(5):853–64.

121. Gosalbes M, Vallès Y, Jiménez-Hernández N, Balle C, Riva P, Miravet-Verde S, et al. High Frequencies of Antibiotic Resistance Genes in Infants’ Meconium and Early Fecal Samples. Journal of developmental origins of health and disease (2016) 7(1):35–44.

122. Schrag SJ, Zywicki S, Farley MM, Reingold AL, Harrison LH, Lefkowitz LB, et al. Group B Streptococcal Disease in the Era of Intrapartum Antibiotic Prophylaxis. New England Journal of Medicine (2000) 342(1):15–20.

123. Klassert TE, Zubiria-Barrera C, Kankel S, Stock M, Neubert R, Lorenzo-Diaz F, et al. Early Bacterial Colonization and Antibiotic Resistance Gene Acquisition in Newborns. Frontiers in Cellular and Infection Microbiology (2020) 10:332.

124. Gonçalves LF, Chaiworapongsa T, Romero R. Intrauterine Infection and Prematurity. Mental retardation and developmental disabilities research reviews (2002) 8(1):3–13.

125. Clayton JB, Vangay P, Huang H, Ward T, Hillmann BM, Al-Ghalith GA, et al. Captivity Humanizes the Primate Microbiome. Proceedings of the National Academy of Sciences (2016) 113(37):10376–81. doi: doi:10.1073/pnas.1521835113.

126. Nagpal R, Wang S, Solberg Woods LC, Seshie O, Chung ST, Shively CA, et al. Comparative Microbiome Signatures and Short-Chain Fatty Acids in Mouse, Rat, Non-Human Primate, and Human Feces. Frontiers in microbiology (2018) 9:2897.

127. Guaraldi F, Salvatori G. Effect of Breast and Formula Feeding on Gut Microbiota Shaping in Newborns. Frontiers in cellular and infection microbiology (2012) 2:94.

128. Chow WL, Lee Y-K. Mucosal Interactions and Gastrointestinal Microbiota. Gastrointestinal microbiology (2006) 1:123–36.

129. Wilson M. Microbial Inhabitants of Humans: Their Ecology and Role in Health and Disease: Cambridge University Press (2005).

130. Miko E, Csaszar A, Bodis J, Kovacs K. The Maternal–Fetal Gut Microbiota Axis: Physiological Changes, Dietary Influence, and Modulation Possibilities. Life (2022) 12(3):424.

131. Ardissone AN, de la Cruz DM, Davis-Richardson AG, Rechcigl KT, Li N, Drew JC, et al. Meconium Microbiome Analysis Identifies Bacteria Correlated with Premature Birth. PloS one (2014) 9(3):e90784.

132. Madan JC, Salari RC, Saxena D, Davidson L, O’Toole GA, Moore JH, et al. Gut Microbial Colonisation in Premature Neonates Predicts Neonatal Sepsis. Archives of Disease in Childhood-Fetal and Neonatal Edition (2012) 97(6):F456–F62.

133. Koren O, Goodrich JK, Cullender TC, Spor A, Laitinen K, Bäckhed HK, et al. Host Remodeling of the Gut Microbiome and Metabolic Changes During Pregnancy. Cell (2012) 150(3):470–80.

134. Sun B, Xu X, Xia Y, Cheng Y, Mao S, Xiang X, et al. Variation of Gut Microbiome in Free-Ranging Female Tibetan Macaques (Macaca Thibetana) across Different Reproductive States. Animals (2020) 11(1):39.

135. Shin N-R, Whon TW, Bae J-W. Proteobacteria: Microbial Signature of Dysbiosis in Gut Microbiota. Trends in biotechnology (2015) 33(9):496–503.

136. Arumugam M, Raes J, Pelletier E, Le Paslier D, Yamada T, Mende DR, et al. Enterotypes of the Human Gut Microbiome. nature (2011) 473(7346):174–80.

137. Chen T, Li Y, Liang J, Li Y, Huang Z. Gut Microbiota of Provisioned and Wild Rhesus Macaques (Macaca Mulatta) Living in a Limestone Forest in Southwest Guangxi, China. MicrobiologyOpen (2020) 9(3):e981.

138. Yasuda K, Oh K, Ren B, Tickle TL, Franzosa EA, Wachtman LM, et al. Biogeography of the Intestinal Mucosal and Lumenal Microbiome in the Rhesus Macaque. Cell host & microbe (2015) 17(3):385–91.

139. Wu Y, Yao Y, Dong M, Xia T, Li D, Xie M, et al. Characterisation of the Gut Microbial Community of Rhesus Macaques in High-Altitude Environments. BMC microbiology (2020) 20:1–16.

140. Adriansjach J, Baum ST, Lefkowitz EJ, Van Der Pol WJ, Buford TW, Colman RJ. Age-Related Differences in the Gut Microbiome of Rhesus Macaques. The Journals of Gerontology: Series A (2020) 75(7):1293–8.

141. David LA, Maurice CF, Carmody RN, Gootenberg DB, Button JE, Wolfe BE, et al. Diet Rapidly and Reproducibly Alters the Human Gut Microbiome. Nature (2014) 505(7484):559–63.

142. Ley RE, Hamady M, Lozupone C, Turnbaugh PJ, Ramey RR, Bircher JS, et al. Evolution of Mammals and Their Gut Microbes. science (2008) 320(5883):1647–51.

143. Chen Z, Yeoh YK, Hui M, Wong PY, Chan MC, Ip M, et al. Diversity of Macaque Microbiota Compared to the Human Counterparts. Scientific reports (2018) 8(1):15573.

