## supplementary Figures and Tables for "Placental and fetal microbiota in rhesus macaque: a case study using metagenomic sequencing"

### Supplementary Materials

#### 1 Supplementary Figures and Tables

##### 1.1 Supplementary Figures

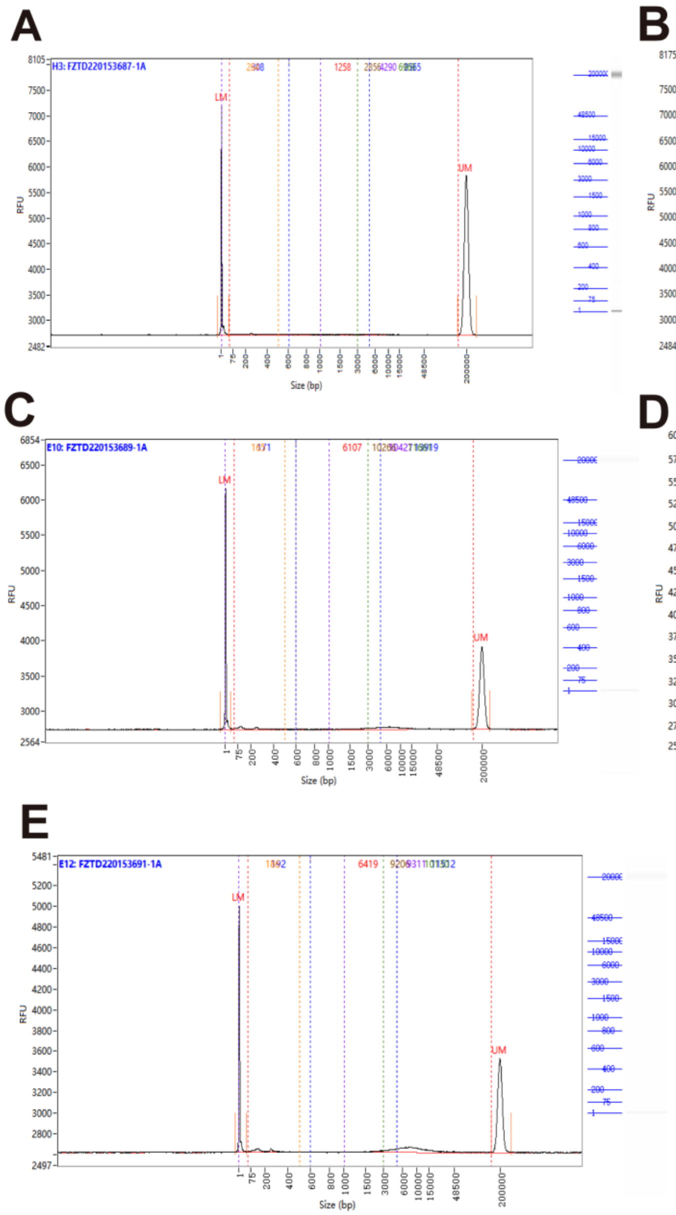

Figure S1 Agarose gel images and AATI fragment analyzer traces. (**A**) Agarose gel image and AATI fragment analyzer trace of workbench environmental swabs. (**B**) Agarose gel image and AATI fragment analyzer trace of operator hand swabs. (**C**) Agarose gel image and AATI fragment analyzer trace of blank swabs. (**D**) Agarose gel image and AATI fragment analyzer trace of blank swabs of PBS solution. (**E**) Agarose gel image and AATI fragment analyzer trace of PBS washing fluid of dissection instruments. Abscissa: fragment distribution of capillary sample where size (bp) reacts; ordinate (RFU) value refers to real-time fluorescence signal intensity of sample during capillary separation process. Higher RFU values indicate higher sample concentrations; Lower marker/Upper marker: refers to reference substance analyzed with sample (not the fragment of the sample itself), used to calibrate the fragment size and concentration of the sample; right side is simulated gel map and corresponding ladder size value label.

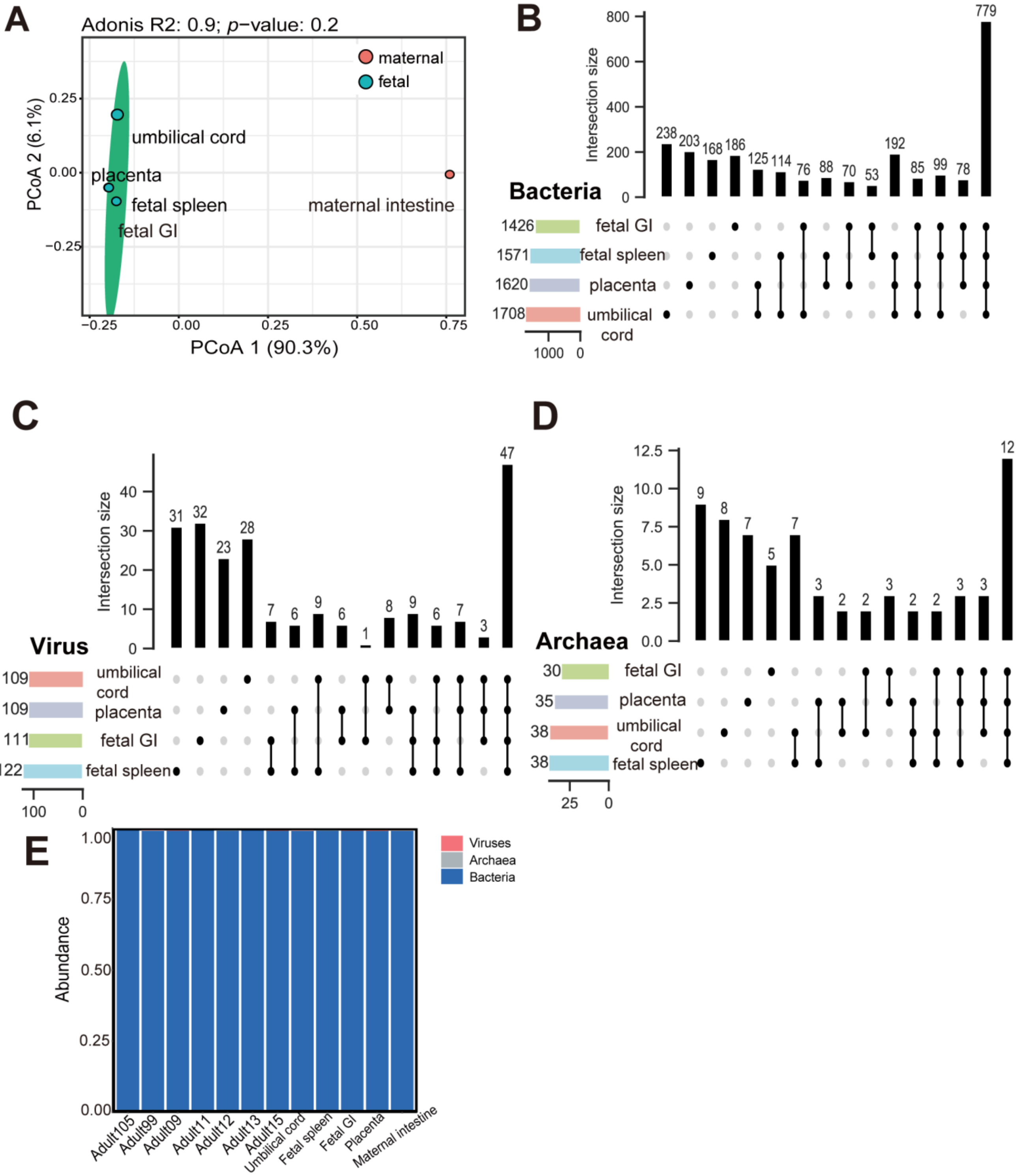


Figure S2 Microbial composition of maternal and fetal samples. (**A**) PCoA plot based on Bray-Curtis distance of species-level relative abundance of microbiota including viruses, archaea, and bacteria between maternal and fetal (Adonis, $R^2 = 0.90$, $p = 0.2$). (**B**) UpSet diagram of exclusive and shared fetal bacteria in different fetal samples. (**C**) UpSet diagram of exclusive and shared fetal viruses in different fetal samples. (**D**) UpSet diagram of exclusive and shared fetal archaea in different fetal samples. (**E**) The proportion of bacteria, viruses and archaea in samples.

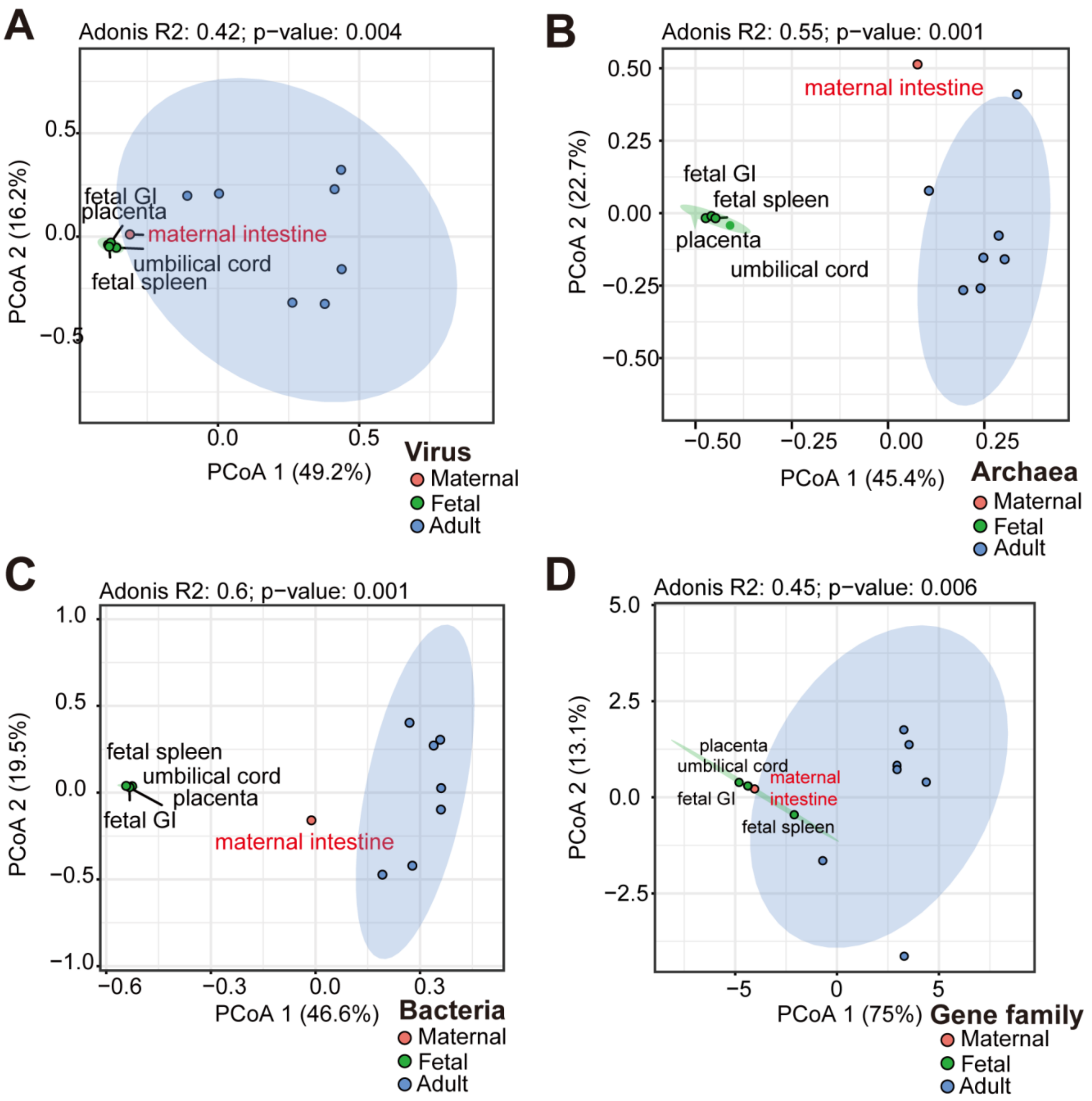


Figure S3 Comparison of microbial diversity and gene families between adult and fetal groups. (**A**) PCoA plot based on Bray-Curtis distance of viruses between two groups (Adonis, $R^2$ = 0.42, $p$ = 0.004). (**B**) PCoA plot based on Bray-Curtis distance of archaea between two groups (Adonis, $R^2$ = 0.55, $p$ = 0.001). (**C**) PCoA plot based on Bray-Curtis distance of bacteria between two group (Adonis, $R^2$ = 0.6, $p$ = 0.001). (**D**) PCoA plot based on abundance of microbial gene families between two group (Adonis, $R^2$ = 0.45, $p$ = 0.006).

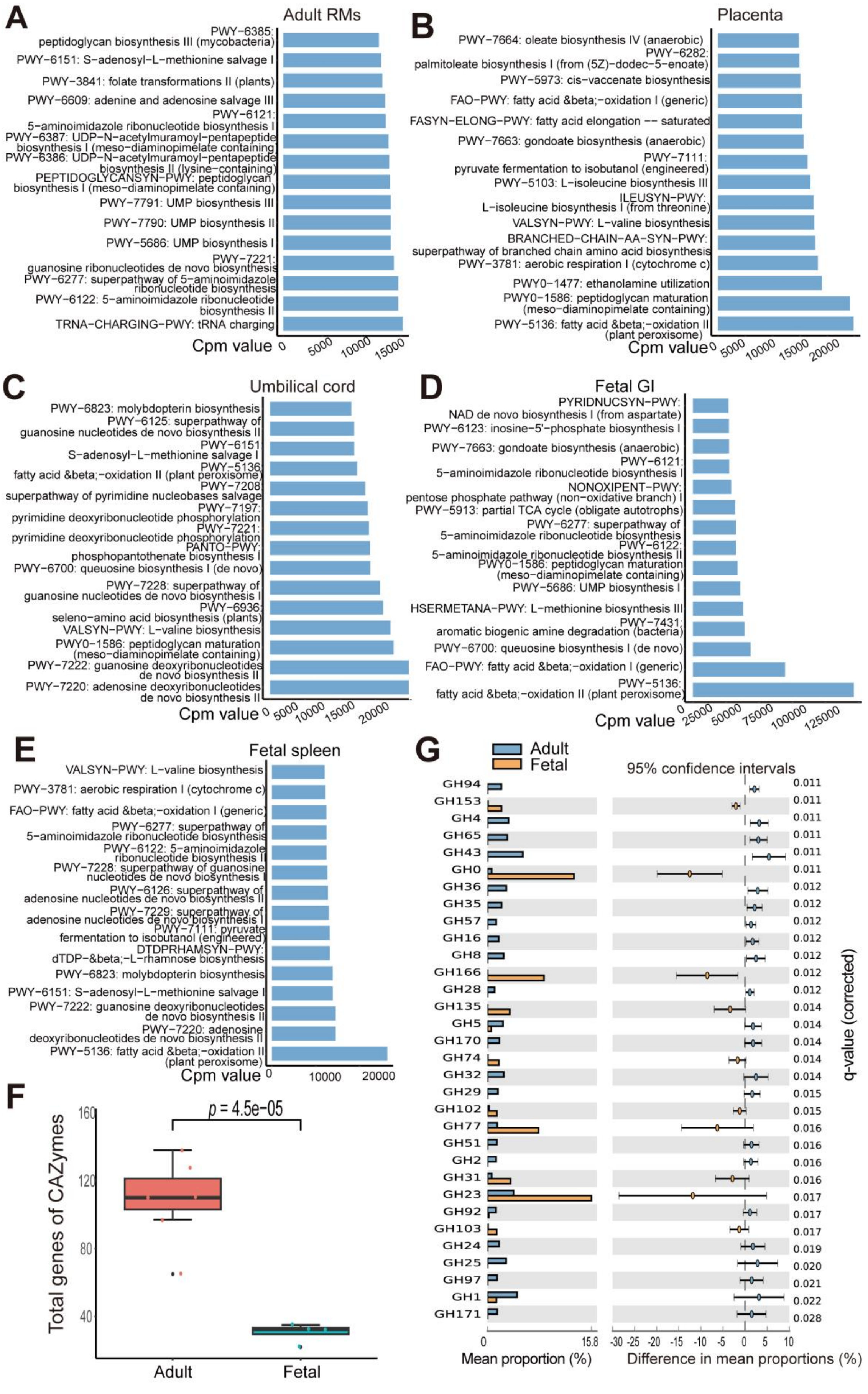
A
Adult RMs
Cpm value
B
Placenta
C
Umbilical cord
D
Fetal GI
E
Fetal spleen
F
Total genes of CAZymes
p = 4.5e−05
Adult
Fetal
G
Adult
Fetal
95% confidence intervals
Mean proportion (%)
Difference in mean proportions (%)
q-value (corrected)

Figure S4 Functional analyses of adult and fetal samples. (**A**) Pathway enrichment analyses of adult gut microbiota. (**B**) Pathway enrichment analyses of placenta microbiota. (**C**) Pathway enrichment analyses of umbilical cord microbiota. (**D**) Pathway enrichment analyses of fetal GI microbiota. (**E**) Pathway enrichment analyses of fetal spleen microbiota. (**F**) Comparison of total genes of CAZymes in adult and fetal microbiomes ($p < 0.05$). (**G**) GH family genes differing in abundance between adult and fetal microbiomes. Bar plot shows mean proportions of GH family genes. Only $p < 0.05$ is shown (Welch's *t*-test, FDR adjusted)

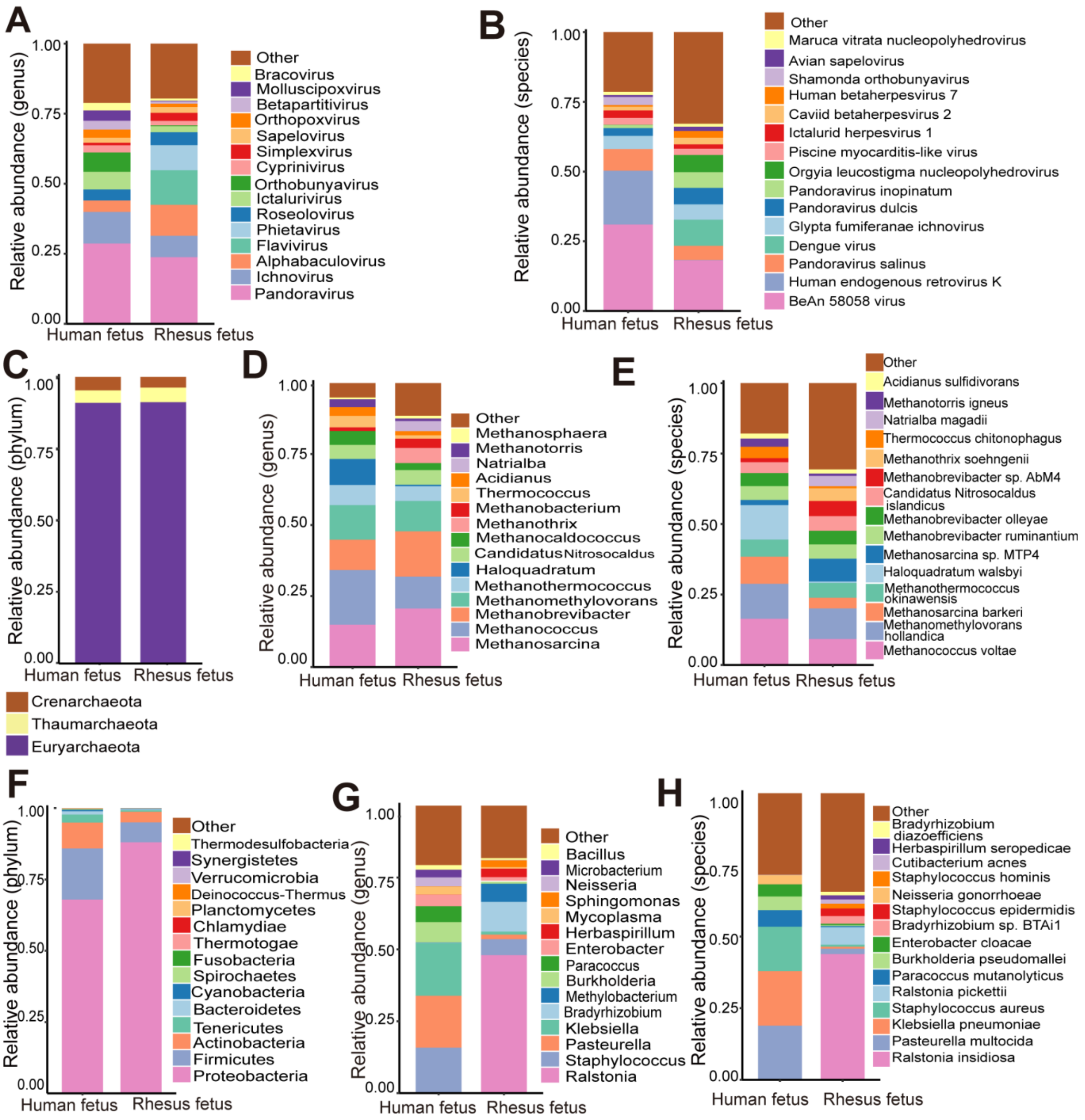


Figure S5 Differential analyses of microbial composition in human and RM fetuses. (**A**) Top 15 abundant virus genera in two groups. (**B**) Top 15 abundant virus species in two groups. (**C**) Top 15 abundant archaea phyla in two groups. (**D**) Top 15 abundant archaea genera in two groups. (**E**) Top 15 abundant archaea species in two groups. (**F**) Top 15 abundant bacteria phyla in two groups. (**G**) Top 15 abundant bacteria genera in two groups. (**H**) Top 15 abundant bacteria species in two groups.

#### 1.2 Supplementary Tables

Table S1 Quality of metagenomic sequencing data.

| Sample | Raw reads | Clean reads | Error (%) | Q20 (%) | Q30 (%) | GC (%) |
|---|---|---|---|---|---|---|
| Fetal GI | 37 300 184 | 36 797 452 | 0.03 | 96.16 | 90.47 | 42.90 |
| Maternal intestine | 37 582 974 | 37 170 980 | 0.03 | 95.55 | 89.08 | 41.51 |
| Fetal spleen | 36 295 682 | 36 006 478 | 0.03 | 96.20 | 90.36 | 41.55 |
| Umbilical cord | 34 152 904 | 33 939 022 | 0.03 | 96.41 | 91.10 | 41.36 |
| Placenta | 34 834 244 | 34 558 488 | 0.03 | 95.97 | 90.09 | 42.60 |

*Error: Average base sequencing error rate; Q20, Q30: Percentage of bases with Phred values greater than 20 and 30 in total bases, respectively; GC: Percentage of GC bases in total bases.

Table S2 Number of shared microbes (including viruses, bacteria, and archaea) between maternal intestine and fetal samples.

| **Type** | **Placenta** | **Umbilical cord** | **Fetal GI** | **Fetal spleen** |
|---|---|---|---|---|
| **Viruses** | 47 | 43 | 50 | 49 |
| **Bacteria** | 587 | 654 | 533 | 590 |
| **Archaea** | 17 | 16 | 15 | 18 |

Table S3 Top five most abundant microbial phyla in RM fetal and adult groups.

| **Level** | **Adult group** | **Fetal group** |
|---|---|---|
| **Bacteria** | Firmicutes | Proteobacteria |
| | Bacteroidetes | Firmicutes |
| | Proteobacteria | Actinobacteria |
| | Actinobacteria | Tenericutes |
| | Spirochaetes | Bacteroidetes |
| **Archaea** | Euryarchaeota | Euryarchaeota |
| | Crenarchaeota | Thaumarchaeota |
| | Thaumarchaeota | Crenarchaeota |

Table S4 Top five most abundant archaea in human and RM fetuses.

| Level | Human fetus | RM fetus |
|---|---|---|
| **Phylum** | Euryarchaeota | Euryarchaeota |
| | Crenarchaeota | Thaumarchaeota |
| | Thaumarchaeota | Crenarchaeota |
| **Genus** | *Methanococcus* | *Methanosarcina* |
| | *Methanosarcina* | *Methanobrevibacter* |
| | *Methanomethylovorans* | *Methanococcus* |
| | *Haloquadratum* | *Methanomethylovorans* |
| | *Methanobrevibacter* | *Methanothrix* |
| **Species** | *Methanococcus voltae* | *Methanomethylovorans hollandica* |
| | *Methanomethylovorans hollandica* | *Methanococcus voltae* |
| | *Haloquadratum walsbyi* | *Methanosarcina sp. MTP4* |
| | *Methanosarcina barkeri* | *Methanobrevibacter sp. AbM4* |
| | *Methanothermococcus okinawensis* | *Methanothermococcus okinawensis* |

Table S5 Top five most abundant bacteria in human and RM fetuses.

| **Level** | **Human fetus** | **RM fetus** |
|---|---|---|
| **Phylum** | Proteobacteria | Proteobacteria |
| | Firmicutes | Firmicutes |
| | Actinobacteria | Actinobacteria |
| | Tenericutes | Tenericutes |
| | Bacteroidetes | Bacteroidetes |
| **Genus** | *Klebsiella* | *Ralstonia* |
| | *Pasteurella* | *Bradyrhizobium* |
| | *Staphylococcus* | *Methylobacterium* |
| | *Burkholderia* | *Staphylococcus* |
| | *Paracoccus* | *Herbaspirillum* |
| **Species** | *Klebsiella pneumoniae* | *Ralstonia insidiosa* |
| | *Pasteurella multocida* | *Ralstonia pickettii* |
| | *Staphylococcus aureus* | *Bradyrhizobium sp. BTAi1* |
| | *Paracoccus mutanolyticus* | *Staphylococcus epidermidis* |
| | *Burkholderia pseudomallei* | *Pasteurella multocida* |
